# Mutagenesis of *orco* impairs foraging but not oviposition in the hawkmoth *Manduca sexta*

**DOI:** 10.1101/541201

**Authors:** Richard A. Fandino, Alexander Haverkamp, Sonja Bisch-Knaden, Jin Zhang, Sascha Bucks, Tu-Anh Nguyen Thi, Achim Werkenthin, Jürgen Ryback, Monika Stengl, Markus Knaden, Bill S. Hansson, Ewald Große-Wilde

## Abstract

Plant volatile detection through olfaction plays a crucial role in insect behaviors. *In vivo*, the odorant receptor co-receptor *orco* is an obligatory component for the function of odorant receptors (ORs), a major receptor family involved in insect olfaction. We used CRISPR-Cas9 targeted mutagenesis to knock-out (KO) *orco* in a neurophysiological model species, the hawkmoth *Manduca sexta. M. sexta* and its host, the Sacred Datura (*Datura wrightii*) share a model insect-plant relationship based on mutualistic and antagonistic life history traits. *D. wrightii* is the innately preferred nectar-source and oviposition host for *M. sexta*. Hence, the hawkmoth is an important pollinator while the *M. sexta* larvae are specialized herbivores of the plant. We generated an *orco* KO through CRISPR-Cas9 to test the consequences of a loss of OR-mediated olfaction in this insect-plant relationship. Neurophysiological characterization revealed severely reduced antennal and antennal lobe responses to representative odorants emitted by *D. wrightii*. In a wind-tunnel setting with a flowering plant, *orco* KO hawkmoths showed disrupted flight orientation and an ablated proboscis extension response to the natural stimulus. However, when testing the oviposition behavior of mated females encountering a non-flowering plant, there was no difference between *orco* KO and wild type females regarding upwind flight orientation and number of eggs laid. Overall, OR-mediated olfaction is essential for foraging and pollination behaviors, but plant-seeking and oviposition behaviors appear largely unaffected.

**Significance statement:** Insects detect plant volatiles mainly through the expression of ORs and IRs on the antennal olfactory sensory neurons (OSNs). In vivo, Orco is an obligate partner for OR, but not IR function and ORs mediate a vast spectrum of olfactory perception. We applied CRISPR-Cas9 in *M. sexta* to mutate the *orco* gene and determine the physiological and behavioral implication of a loss of Orco receptor function in a semi-ecological interaction with D. wrightii. We found that while behaviors related to foraging were largely disrupted, other sensory modalities outside Orco function determine the relationship between an ovipositing female and its plant host. These results have implications toward understanding the olfactory basis of insect-plant interactions shaping our ecological and agricultural landscapes.

## Introduction

Chemical signals are important for feeding and reproductive behaviors in insects and are major drivers of insect-plant interactions (1, 2). In the North American southwest, the Sacred Datura *Datura wrightii* has a relationship with the hawkmoth *Manduca sexta* based on pollination and herbivory (3–5). Large and light reflective *D. wrightii* flowers advertise nectar through a fragrant volatile blend and are an innately preferred nectar source for foraging *M. sexta* (5, 6). Both visual and olfactory cues of the upright trumpet flower guides foraging decisions, indicated by hovering and an unfurling proboscis (7, 8). The hovering hawkmoth with extended proboscis is capable of close range olfactory (9) and mechanosensory (10) guidance to reach the nectar canals. Field observations find that females intersperse feeding bouts with oviposition (11) and *M. sexta* females demonstrate innate attraction not only to flowering but also to non-flowering *D. wrightii* plants (12). Plant volatiles elicit orientation and directed flight in the female hawkmoth (12–14), followed by tarsal contact and abdomen curling behavior, likely recognizing contact chemosensory cues eliciting oviposition (15). Plants are also capable of manipulating their environment through modulation of the volatile blend. For example, herbivore - induced volatiles may facilitate attraction of the natural enemies of the herbivore (16–18) or repel a gravid female (19–21). Therefore, the detection of plant volatiles poses a critical factor balancing the survival of insect and host through olfactory mediated behaviors.

*D. wrightii*, like most plants, produce a large repertoire of volatile compounds and the *M. sexta* antennae respond to many of these compounds (14, 22–24). Instead of perceiving the plant as a whole, *M. sexta* distinguishes the olfactory identity of its host through a behaviorally relevant odorant or odorant blend. For example, a minimal blend of three odorants representative of the full floral volatile mediate odor modulated upwind flight (25, 26). Subsequently, close range feeding and oviposition behaviors are elicited by single compounds (27). The hawkmoth is able to generalize the olfactory stimulus of the plant through a spatiotemporal neural activation pattern elicited by a particular odorant blend or single compound in the first processing center of the insect brain, the antennal lobe (AL) (25–27). Activation patterns of the AL convey a message to higher olfactory centers of the brain eliciting a behavioral output. However, information from other sensory modalities may be integrated with olfactory information through multimodal mushroom body output neurons in these higher order brain centers (28, 29). Experimental set-ups in the wind-tunnel often set a minimalized visual target and olfactory stimulus to measure feeding or oviposition behaviors, however insect behaviors may be highly adaptive when confronted with natural multi-sensory stimuli (30–32), making the interpretation of a single sensory system in an ecological context difficult.

The main insect olfactory sense structures are the antennae and maxillary/labial palps. Here, odorants are detected by chemosensory receptors belonging to protein families of gustatory receptors (GRs), ionotropic receptors (IRs), and odorant receptors (ORs) (33, 34). GRs, IRs, and ORs recognize specific molecules or conditions in the environment, resulting in the activation of associated olfactory sensory neurons (OSNs), which send axonal projections to the AL. GRs are largely involved in contact chemoreception, for example in the proboscis and tarsi, where in Lepidoptera they play an important role in feeding and oviposition behaviors (35). Besides the detection of non-volatile chemicals, GRs found in insect antennae or palps detect CO_2_ (31, 36), which is a constituent of the *D. wrighitii* flower volatile blend (37). IRs are attributed to physiological responses to volatile acids, amines, and aldehydes, but are also involved in other sensory modalities, like thermo- and hygroreception (38–40). The seven transmembrane ORs are involved in the majority of odorant binding and detection. They colocalize with the conserved odorant Co-Receptor (Orco), which serves as obligatory chaperone for the localization and maintenance of ORs in the OSN membrane cilia (41, 42). On its own, Orco forms a homotetramer with cation channel function (43–47). The ability for the receptor channel to open during the neuronal resting potential contributes a pacemaker channel function controlling spontaneous activity of the OSN (48, 49). The precise role of the heteromeric ORCO-OR receptor complex in odorant signal transduction remains under debate (46, 50–53). However, there is consensus that the Orco-OR receptor complex binds a cognate ligand which, finally, initiates ion currents that result in depolarization of the sensory neuron. While ORs are highly diverse, e.g. there are 73 transcriptionally active ORs in both male and female *M. sexta* (54), there is only one evolutionarily highly conserved Orco (55–57).

The *orco* null mutations within different insect orders are associated with a critical olfactory loss of function towards a broad range of odorants (42, 58–63). Additionally, olfactory related behaviors such as short distance orientation (42), host-chemical cue detection (58), mating (59, 61), and social communication (62, 63) are disrupted. We examined the role of OR-mediated olfaction within the *M. sexta* – *D. wrightii* interaction. We developed a specialized microinjection protocol for delivery of CRISPR-Cas9 ribonucleoprotein (RNP) complex to *M. sexta* embryos and generated an *orco* KO hawkmoth. The established *orco* null mutant, which was confirmed through genotyping, lost pheromone detection and the male *orco* Kos did not copulate (59, 61). The *orco* KO individuals showed a deficiency in response to a selected panel of odorants largely representative of *D. wrighitii* associated volatiles. We used the established *orco* KO line to test the contribution of OR-dependent olfaction in foraging and oviposition behaviors using the natural multi-sensory stimulus of a *D. wrightii* plant. Our study offers the first analysis of how ablation of OR-dependent olfaction affects foraging and oviposition behaviors of insects in relation to the plant and the extent to which distinct chemosensory cues guide both behaviors. The results provide novel insights into the integration of sensory mechanisms determining both mutualistic and antagonistic interactions between insects and plants.

## Results

### CRISPR-Cas9 generation of *M. sexta orco* null mutation

We performed CRISPR-Cas9 experiments to mutate orco and obtain a non-functional Orco protein. The *M.* sexta Orco protein has 7 transmembrane regions and a molecular weight of ∼ 54kDa (49) (Fig. 1A). A 10 - exon gene (Msex2.12779) in the official gene set 2 (OGS2) is predicted to encode the *M. sexta orco* gene (Fig. 1B *top*). It is located on the reverse strand of the JH668978.1 genomic scaffold. We used the CHOPCHOP v2 web tool on the OGS2 sequence data (64, 65) to identify CRISPR single guide RNA (sgRNA) sequences against the second exon of the *MsexOrco*, based on reduced likelihood of off-target sites in the *M. sexta* genome. From multiple sgRNA candidates, we selected two that were situated on opposite strands of the second exon, 51 nucleotides apart from the respective CRISPR – Cas9 PAM sites (Fig. 1B). A ribonucleoprotein (RNP) consisting of multiplexed sgRNAs in complex with the Cas9 enzyme were injected into 1 – 2 hour old eggs. We excised the second to third instar caterpillar horn tissue from surviving caterpillars (n = 42) of 126 RNP injected individuals. Using the T7 Endonuclease 1 (T7E1) assay on PCR product we were able to discern insertion deletion mutations based on cleavage of heteroduplexed double stranded DNA (Fig. S1B). Of all injected G0 caterpillars (n = 42), 95% contained *MsexOrco* mutations. G0 individuals were backcrossed with wild type (WT) hawkmoths, allowing isolation and sequencing of germ-line mutations (Fig. S1A). Inherited mutations were identified using T7E1 on the PCR product of G1 colonies (Fig. S1C). Three mutations from three separate parental pairs were identified by T7E1 digest of PCR product, cloning (TOPO-TA) and sequencing of positive individuals (Fig. 1B *bottom*, Fig. S1C). Mutation 1 was chosen based on both likelihood of functional consequences and feasible PCR based genotyping (Fig. 1C). For this study, we chose an 11 bp insertion combined with a 1 bp deletion (Fig. 1B, C *top*). The 11 bp with 1 bp deletion generated a frameshift with 2 premature stop signals on the predicted first extracellular loop (Fig. 1A, C bottom).

**Fig. 1.**
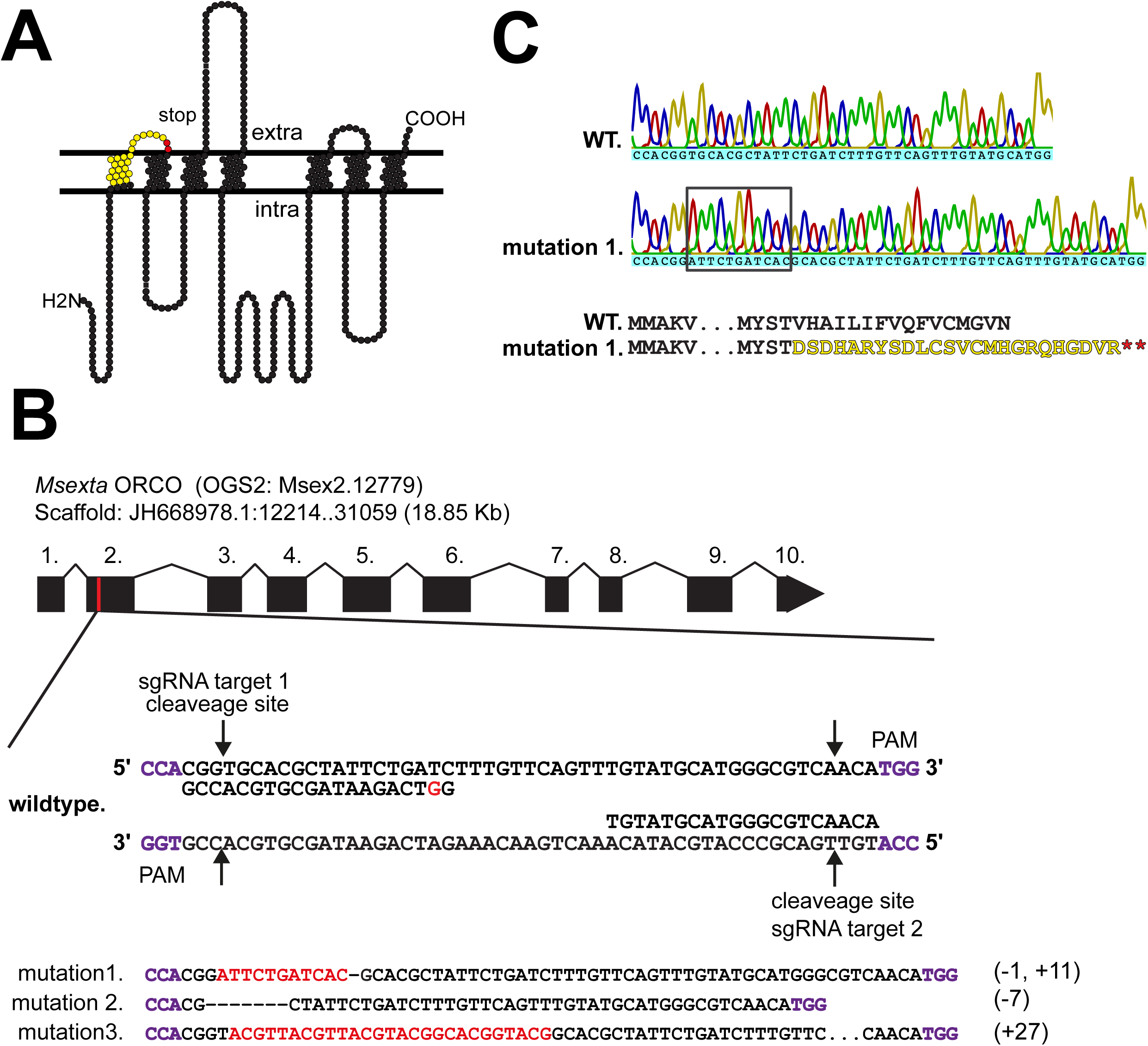
The *MsexOrco* gene was mutated using CRISPR-Cas9. (A) The *M. sexta* Orco protein has 7-transmembrane domains. The first transmembrane domain was mutated generating a frameshift (yellow) and translational stop signals (red) (B) *Top*. The *M. sexta* 10 - exon *orco* gene (18.85 Kb) was accessed through the annotated official gene set (OGS) (gene id Msex2.12779), with scaffold and base pair coordinate location on the positive strand (12,214 – 31,059 bp) within scaffold JH668978.1. A proximal region to the start of exon 2 was targeted for CRISPR single guide RNA (sgRNA) design. Two sgRNAs (target1 and target2) on opposite DNA strands were multiplexed and injected. Target 1 was designed with a mismatch nucleotide, indicated in red to raise in vitro transcription efficiency. PAM sites indicated in purple, Cas9 cleavage sites 3 nt from the PAM site indicated by arrows. *Bottom*. Three germ-line insertion deletion (indel) mutations were recovered and sequenced (mutation 1, 2, 3). All indels were generated by sgRNA target 1, insertion indicated in red, deletions indicated as dash lines. (C) *Top*. Mutation 1 was used for downstream functional analysis. Sanger sequencing chromatogram of wildtype (WT) individual and mutation 1 allele shows insertion event, indicated by box. *Bottom*. Indel generated a frameshift mutation introducing two stop codons downstream, giving rise to a truncated 72 residue protein, indicated through the predicted amino acid sequence.

### Electrophysiological confirmation of loss of pheromone olfaction in genotyped *orco* KO

A BclI restriction enzyme cutting site was introduced in the mutation1, not found in the WT allele, facilitating the identification of the KO allele. The individuals with the orco KO allele produced a 160 and 52 bp product, no cuts were noticed for the WT individuals (Fig 2A). To test for functional consequences of the chosen *M. sexta orco* mutation, we analyzed antennal and brain responses to the sex pheromone of genotyped individuals. We tested the whole excised antennal response using electro-antennograms (EAG). Male WT and *orco* heterozygous (HET) antennae exhibited significant responses of ∼ 2 mV from bombykal stimulation (Fig. 2B *left*), while *orco* KO responses to the pheromone were ablated (Fig. 2B right). The long trichoid sensilla of the *M. sexta* male antennae house two OSNs, with one responding to the main female pheromone component (E,Z)-10,12-hexadecadienal (bombykal) (22, 66) that is detected by the pheromone receptor MsexOR-1 (51). We employed single sensillum recordings (SSR) of pheromone-sensitive trichoid sensilla in the homozygous *orco* KO genotype, compared to WT, and HET hawkmoths in search for a loss of function of the bombykal sensing OSNs. The SSR recordings from WT and *orco* HET hawkmoths found highly significant responses after stimulation with bombykal, while the *orco* KO did not show any significant response (Fig. 2C *right*). Next, we investigated the neurophysiology of *orco* KO animals by recording the activity of the antennal lobe (AL). The AL receives projections from the antennal OSNs, and is the first olfactory processing center of the insect brain. Using fluorescence imaging, we visualized calcium-dependent activation patterns in the AL of WT (n = 4) and *orco* HET males (n =4) in response to stimulation with bombykal. We found strong neural activity at the entrance region of the antennal nerve, where the pheromone processing macro-glomerular complex (MGC) is located (67). In contrast, no such neural activity could be observed in the AL of *orco* KO males (n = 4) (Fig. 2D). Also, a morphological analysis of the male AL showed that the MGC, which is innervated by pheromone sensitive *orco* expressing OSNs, had a smaller relative volume in *orco* KOs in comparison to WTs (Fig. S2). These results, along with the inability of male *orco* KOs to copulate indicated that the *orc*o mutation genotyped through Bcl1 restriction digest indeed led to a loss of function phenotype. OR-mediated pheromone detection was absent in the *orco* KO male hawkmoths. Thus, after genotyping each experimental animal, EAG recordings were employed in all adults stimulating with pheromones and other odorants, to further confirm the accuracy of genotying performed in all caterpillars (n_male_ = 88; n_female_ = 191).

**Fig. 2.**
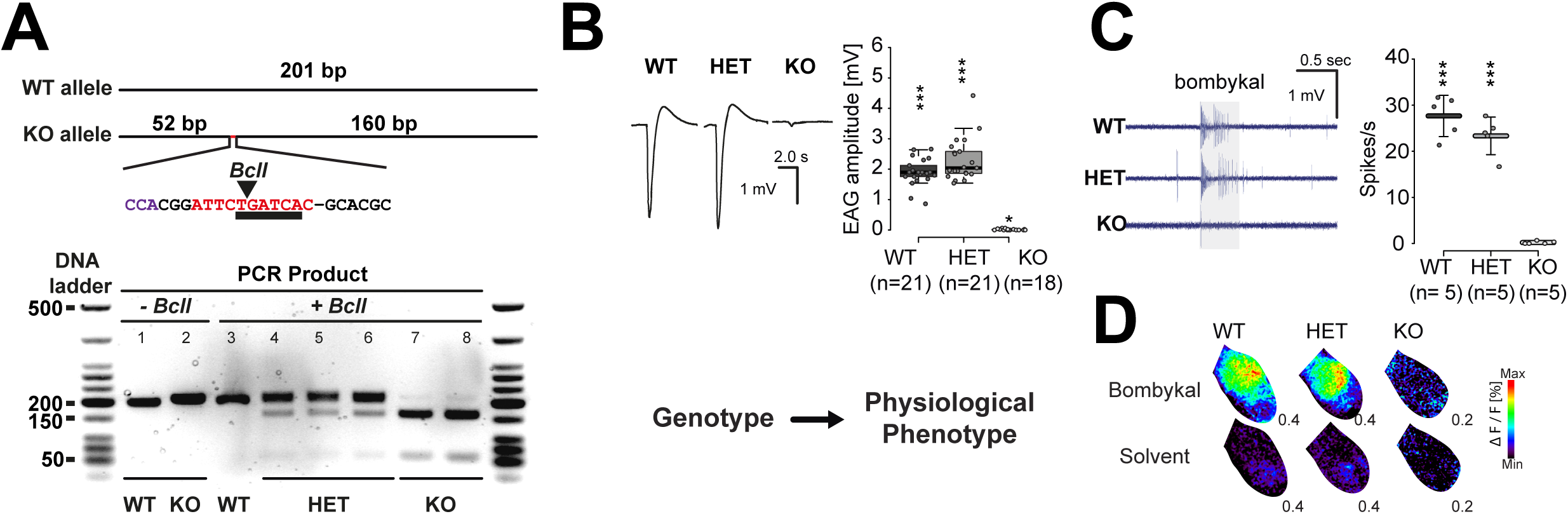
Genotyped *M. sexta orco* KOs do not respond to pheromone in male antenna and antennal lobe (AL). (A) *Top.* Primers designed to amplify 201 bp of the targeted exon 2 of *M. sexta* orco. Knock-out (KO) allele produces a 212 bp amplicon, characterized by the (−1, +11) indel introducing a Bcl1 restriction enzyme cutting site not found in the WT allele. Insertion indicated in red, restriction site underlined arrow points to enzymatic cut site. *Bottom*. PCR product amplified from individual caterpillar horn tissue samples loaded on a 2.0 % agarose gel. (Lanes 1 and 2) PCR product with no restriction enzyme added, (Lanes 3-8) PCR product with restriction enzyme. Lane 3, no cut bands identify a WT caterpillar. Lanes 4-6, ∼ 200 bp band of PCR product is still dark, with lighter bottom bands at ∼ 150 and 50 bp, indicative of heterozygous (HET) individuals. Lanes 7 and 8, ∼ 200 bp is mostly digested and breaks down to a darker ∼ 150 bp band and a slightly darker ∼ 50 bp band, indicative of the homozygous KO individual. (B) EAGs from clipped male antennae stimulated with bombykal; same concentration as in (C and D). *Left*, representative EAG recordings; *right*, boxplots show the net response of n = 21 / WT; n = 21 / HET; n = 18 / KO; *, p ≤ 0.05, ***, p ≤ 0.001, Wilcoxon rank sum test *versus* zero). Boxplots show the median net EAG amplitude (horizontal line in the box), the 25^th^ and 75^th^ percentiles (lower and upper margins of the box) together with the 1.5 x interquartile range (whiskers), and individual data points (circles). Genotyped individuals were all confirmed with EAG recordings, matching genotype to a physiological phenotype. (C) In single sensillum recordings (SSR) from long trichoid sensilla of adult male antennae were stimulated with 10^-2^ (v/v) bombykal in mineral oil for 0.5 ms. *Left*, representative SSR traces, odor stimulation indicated by grey bar; *right*, mean net responses (± SD), circles represent results from individual animals (n = 5 moths / genotype; ***, p ≤ 0.001, n.s., p ≥ 0.05, one-sample t-test *versus* zero). (D) Neural activity patterns in the male AL after stimulation with bombykal (*top panel*) or solvent (*bottom panel*). False color-coded images are representative Ca^2+^ imaging recordings from right ALs, the entrance of the antennal nerve is in the upper left; images were generated by subtracting the frame before stimulus onset from the frame with the maximum response. Calcium activity was normalized for each image and color-coded (see color bar); the maximum activity image frame (arbitrary scale) is given at the bottom right of each image.

### *M. sexta orco* KOs showed disrupted responses towards *D. wrightii* volatiles

After establishing that the selected mutation selected consistently resulted in loss of pheromone detection, we investigated the responses to host plant odors. To determine, wheteher, *M. sexta* orco KOs still detect *D. wrightii* odors we analyzed antennal and AL responses to stimulation with headspace collected from flowering plants (Fig. S3). We found extensive calcium activation patterns throughout the AL of both male and female WT and *orco* HET hawkmoths, but no activity in the AL of KO hawkmoths (Fig. 3A, *top*). Antennae of both sexes of WT and *orco* HET genotypes showed responses that were significantly higher than to stimulation with solvent alone. However, no responses were detected in *orco* KO hawkmoths (Fig. 3A, *bottom*). Next, we tested common floral and leaf volatile compounds as well as amines and acids to cover also the typical response spectra of IRs (68). We included five odorants that are characteristic of *D. wrightii* floral odor bouquet. These were the three behaviorally relevant floral compounds linalool, benzyl alcohol and benzaldehyde, which in combination elicit directed flight and foraging behavior (25) and the monoterpenes, geraniol and eucalyptol (69). The antennal responses to most compounds in both male and female *orco* KO hawkmoths were strongly reduced or absent (Fig. 3B, Fig. S4, S5, Table S1, S2). Out of the three behaviorally sufficient floral compounds, benzaldehyde still elicited responses from both male and female *orco* KO antennae. When we analyzed the activation pattern in the *orco* KO AL for benzaldehyde, we found that although most of the activity elicited in WT and *orco* HET hawkmoths had been lost, populations of neurons in a limited region opposite to the entrance of the antennal nerve were still activated (Fig. 3C). Glomeruli in this region were described to mainly respond to amines, acids, and aldehydes (27), a response spectrum that is characteristic for IRs, but not for ORs (68). When testing odorants associated with *D. wrightii* leaves the behaviorally active green leaf volatiles (GLVs) trans −3 and cis - 2 hexenyl acetate evoked stronger responses than the sesquiterpenes, farnesene, beta-caryophyllene and alpha-humulene (Fig. 3B). None of these odorants were predominantly active in either male or female *orc*o KO (Fig. 3B, S4, S5, Table S1, S2). Notably, among the tested compounds, hexanoic acid, which is found in different host plants of *M. sexta* (12), caused the strongest antennal activity in female *orco* KO antennae, although the response was still weaker than in WT or *orco* HET antennae. The calcium activity in the orco KO AL evoked by hexanoic acid was not as widespread as in the WT or *orco* HET hawkmoths but limited to a distinct part of the AL (Fig. 3C) that was similar to the region activated by benzaldehyde. The majority of compounds that activate IR-expressing neuron in the antennae of *D. melanogaster* are amines or carboxylic acids (40). Therefore, we tested the *M. sexta* antennae for carboxylic acids and two amine compounds. Isovaleric acid also elicited pronounced responses in WT and *orco* HET and a significant response in the *orco* KO, but the other tested acids and amines did not elicit clear responses in WT, *orco* HET or *orco* KO (Fig. 3B, S4, S5, Table S1 and S2).

**Fig. 3.**
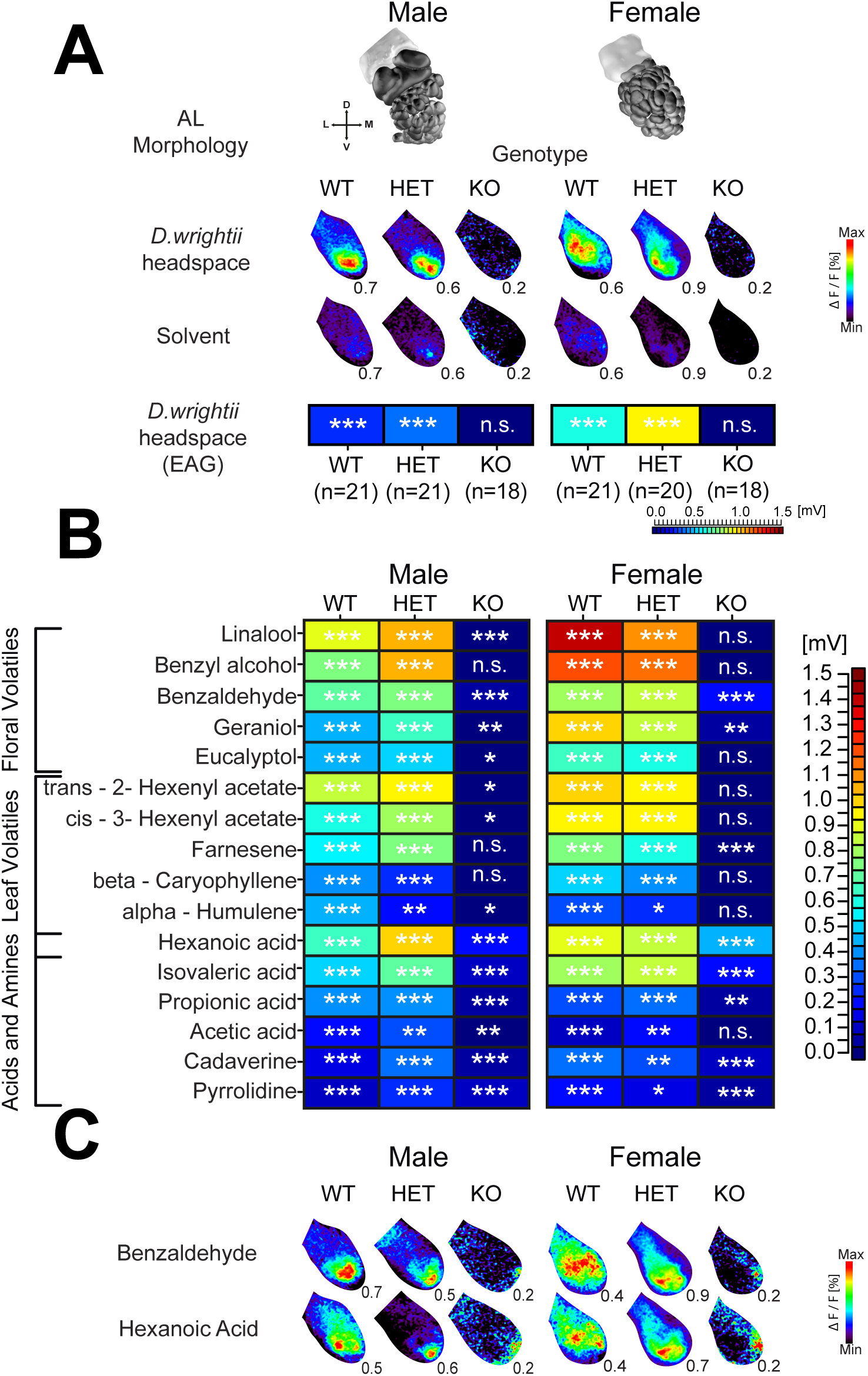
*M. sexta orco* KOs show disrupted or reduced olfactory responses to both single ecologically relevant odors and to whole plant headspace. (A) *D. wrightii* headspace collection (SuperQ), solved in dichloromethane (DCM) was used to test whole excised male and female antennal response (for gas-chromatogram see Fig. S1). *Top.* Neural activity patterns in male and female AL after stimulation with D. wrightii headspace in DCM (solvent). Pattern of response varies based on sexual dimorphic morphology of the *M. sexta* AL (*top*, Grosse-Wilde et al., 2011). *Bottom*. Heat map representation of median EAG response values (mV) corrected for the response to solvent from male and female antennae; ***, p ≤ 0.001, n.s., p ≥ 0.05, Holm corrected Wilcoxon rank sum test versus zero. (B) Male and female EAG responses towards single odorants. The sample size is the same as (A); (*** p ≤ 0.001, ** p ≤ 0.01, * p ≤ 0.05, n.s not significant from 0; Wilcoxin rank sum test versus zero). Box plot representation and individual P-values are shown in Fig. S4, S5 and table S1, 2. Medians were converted to heat map color-scheme (see color bar). (C) Neural activity in the AL of males and females after stimulation with benzaldehyde (floral odor) and hexanoic acid (plant) as the strongest EAG eliciting odorants, in the *orco* KO, among their respective sets.

### Disrupted host localization and foraging behaviors in *orco* KO

The mutualistic relationship between *M. sexta* and *D. wrightii* has been a focus of study in natural and laboratory settings, leading to various insights into the behavioral and sensory mechanisms involved in this interaction (3, 11, 70, 71). After characterizing *orco* mediated olfactory deficits in the *orco* KO to headspace and representative odorants from *D. wrightii,* we tested how these deficits would affect the hawkmoth foraging behavior. We placed a *D. wrightii* plant with a single flower in a wind tunnel and released a naive hawkmoth approximately 220 cm downwind. Regardless of genotype nearly all tested male hawkmoths approached the flowering plant (Fig. 4A, C, for similar results from virgin females see Fig. S6) i.e. they flew upwind and either hovered in front of the flower with an unfurled proboscis, or simply alighted on the plant. *Orco* KO hawkmoths took longer from takeoff to first contact with the flower compared to both control genotypes (Fig. 4B, S7). These differences were due to anemotactic behaviors of the WT and *orco* HET hawkmoths in comparison to stochastic flight of *orco* KO hawkmoths (Fig. 4C, C’). When approaching the *D. wrightii* flower almost all WT and *orco* HET hawkmoths unfurled their proboscis and probed the outer petals and the inner corolla (Fig. 4D, E, S8, Supp. Video S1).

**Fig. 4.**
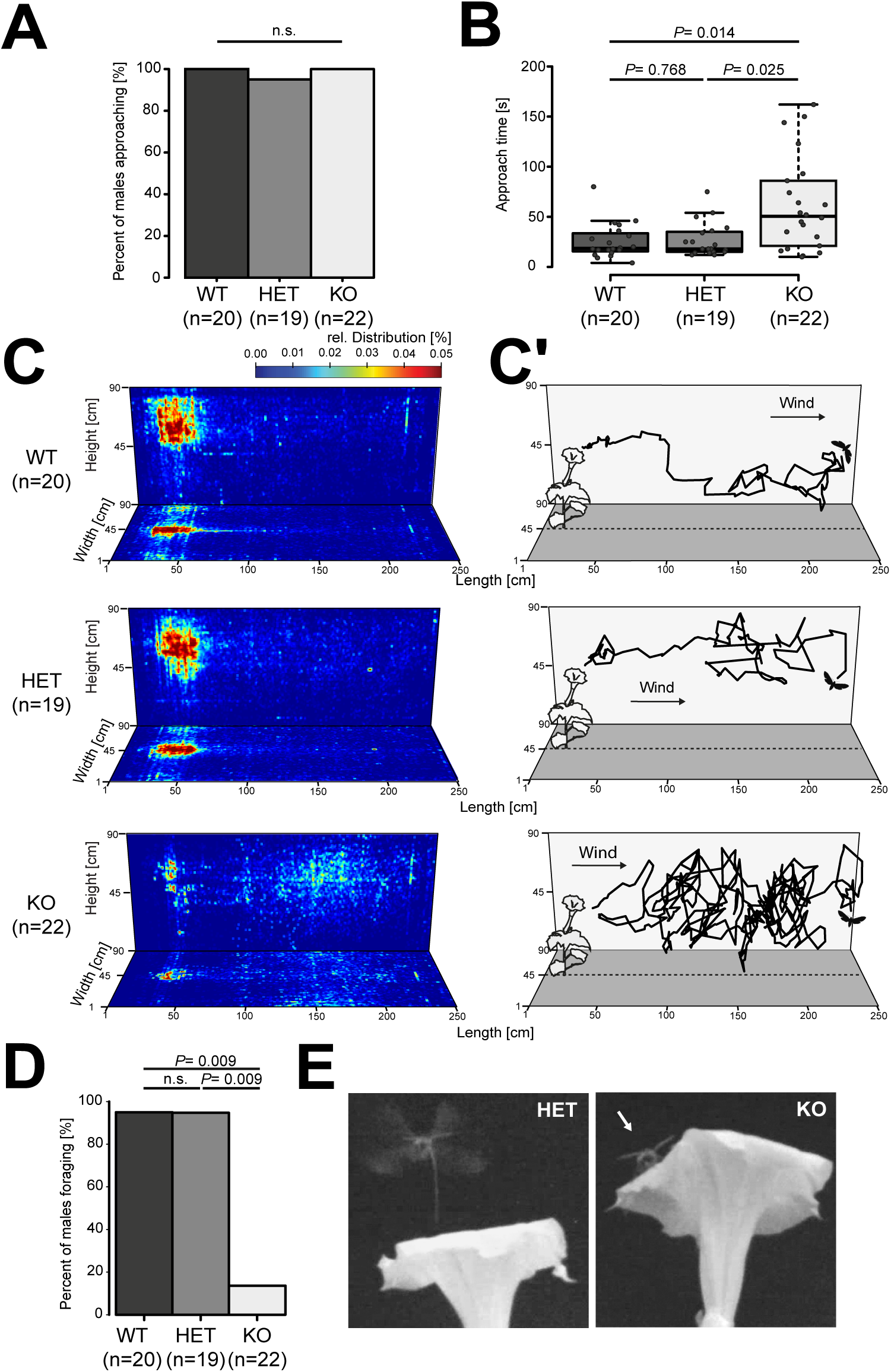
*M. sexta orco* KO took longer to reach flowering plant and showed disrupted proboscis extension behavior. (A) Percentage of male hawkmoths that flew towards a flowering *D. wrightii* plant (stimulus source) in a 2.5 m long wind tunnel. There was no significant difference (n.s.) between genotypes (Holm corrected Fisher’s exact test, two-sided). (B) Approach time was calculated from take-off until individuals hovered in front of the flower or contacted the plant in a 5-minute free flight experiment. Box plots indicate median with 25th and 75th quartile of the data with 1.5x the interquartile range (whiskers), filled circles representing individual data points. P-values based on Holm corrected Wilcoxon rank sum test. (C) Heat map representation of relative distribution of all summed flight tracks per genotype. Single pixels represent points of relative transit of the hawkmoth color coded from maximum transit (red), no transit (dark blue) (see color bar). (C’) Representative individual flight track schematic, hawkmoth (not to scale) depicts start of flight towards flowering plant (not to scale). (D) Percentage of approaching males that foraged. Foraging counted when males unfurled their proboscis and contacted the flower with proboscis (Holm adjusted Fisher’s exact test). (E) Image capture from an infrared filter camera above the flower in wind tunnel flight experiment. *Orco* HET individual with proboscis extended getting ready to forage and *orco* KO individual resting on the flower, without proboscis extended.

Furthermore, the hawkmoth with extended proboscis, often crawled in and out of the flower to feed on different nectar chambers within the *D. wrightii* flower. In contrast, the majority of *orco* KO hawkmoths did not extend their proboscis, but abruptly alighted on the flower or plant (Fig. 4D, E, S8). Only four out of 31 tested male and female *orco* KO individuals probed the flower with their proboscis, three of which attempted to feed. This likely indicates that although the *orco* KO hawkmoths were able to search, they were unable to identify the flower as a nectar source, with disrupted olfaction.

### Host localization and oviposition were not significantly altered in the mated female *orco* KO

Host localization by a gravid female is the initial step in the antagonistic interaction between *M. sexta* and *D. wrightii.* After arriving at a plant, the female hawkmoth holds on to the leaf with her forelegs, curls her abdomen and holds this position in a characteristic motion (27), possibly to assess plant quality through chemoreceptors on the ovipositor (72). When a suitable oviposition site has been identified, the hawkmoth places a single egg and moves to another position on the plant. We exposed mated female hawkmoths to a non-flowering *D. wrightii* plant in a wind tunnel. Independent of the genotype, the majority of mated females contacted the plant (Fig. 5A). Although the time to contact was not significantly different between genotypes (Fig. 4B), the *orco* KO animals seemed to approach the plant in a less directed flight pattern (Fig. 5C, C’). All three *M. sexta* genotypes investigated the whole plant after contact (Fig. 5C). Nearly all mated females exhibited oviposition and egg laying behavior after contacting the plant, with no differences between the three genotypes (Fig. 5D, Supp. Video S2). Additionally, out of the females that made contact, all three genotypes placed a similar number of fertile eggs on the plant (Fig. 5E); indicating contact as the more crucial sensory stimulus in the oviposition response after localizing the host.

**Fig. 5.**
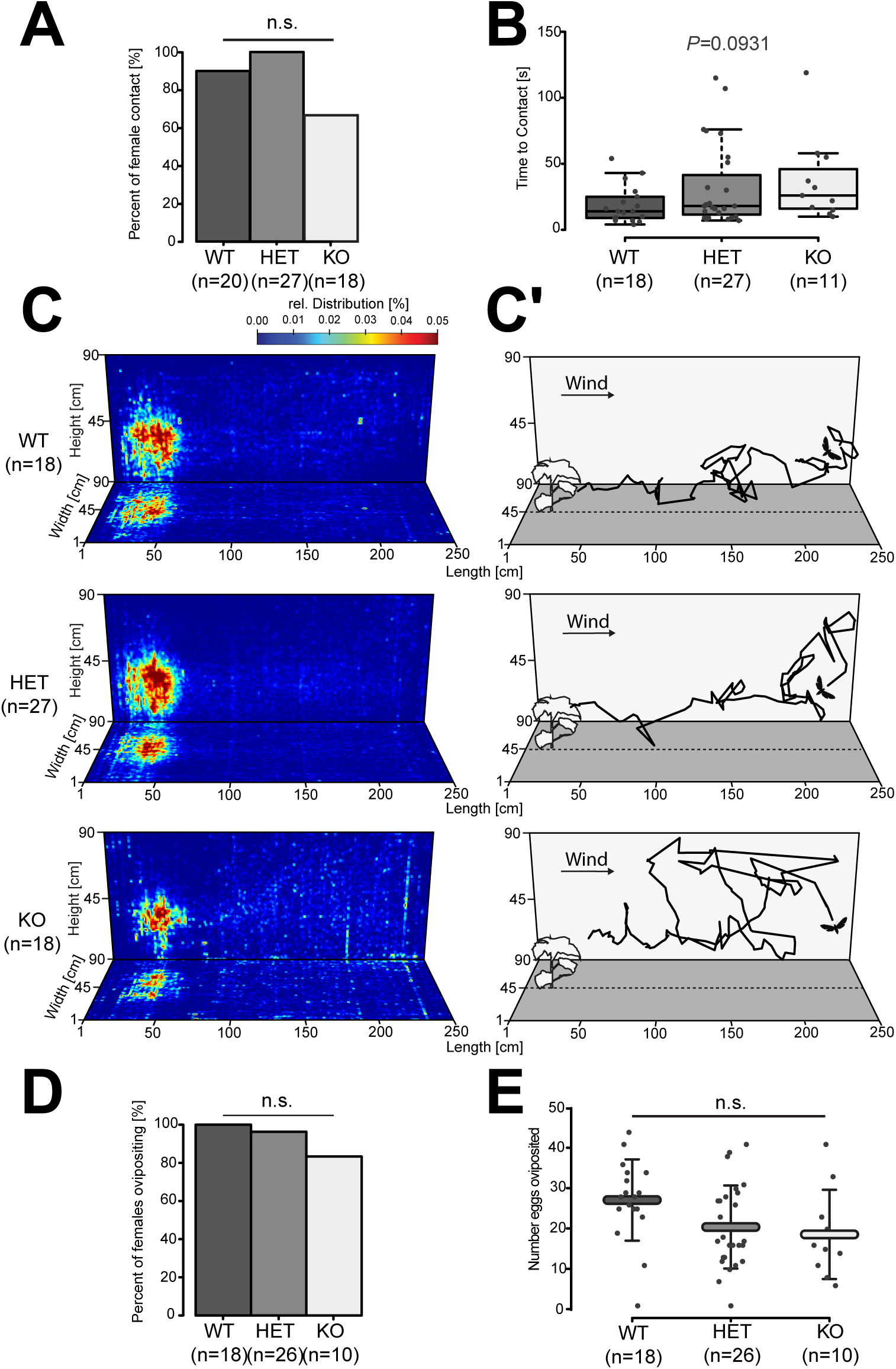
Gravid *orco* KO females landed and oviposited on *D. wrightii* plant. (A) Percent of mated *M. sexta* females that flew towards a non-flowering *D. wrightii* plant was not significantly different between genotypes (n.s.) (Holm adjusted Fisher’s exact test). (B) Time for individual hawkmoths to approach and make contact with the plant was recorded within a 5-minute trial. Boxplots indicate the median and 25th and 75th quartile of the data as well as 1.5x interquartile range (whiskers), points denote individual measurements. There was no significant difference between genotypes (n.s.), Kruskal-Wallis H-test. (C) Heat map representation of relative distribution of all mated female hawkmoths per genotype in a wind tunnel, heat map values represent summed times of all individual flights, pixel regions represent a hawkmoth relative transit point. (C’) Representative flight traces of a single individual. (D) Percent of individual mated females that approached and oviposited (contact on leaf and abdomen curl) on whole *D. wrightii* plant were not significantly different between genotypes (n.s.; Holm-corrected Fisher’s exact test). (E) Total number of eggs laid on *D. wrightii* plants during 5-minute wind-tunnel flight trial. Bars and error bars represent mean ± SD, with no significant differences between genotypes (n.s.; One-way ANOVA).

## Discussion

In an ecological context other sensory cues often accompany olfactory signals, making it challenging to determine the relative importance of these signals for a certain behavioral task. We addressed this by generating an *orco* KO hawkmoth with CRISPR-Cas9 and identified a largely ablated response to *D. wrightii* headspace and representative odorants of floral and foliage volatiles. The orco-KO background allowed us to quantify and qualify a response towards certain acids and aldehydes. Antennal responses towards these classes of odorants were reduced but not completely ablated. The highly disrupted olfactory response allowed us to observe a critical function for OR-mediated olfaction in flight and proboscis extension behaviors associated with foraging. In contrast, oviposition behaviors were functional without OR-mediated olfaction.

We were able to highlight two distinct contextual uses of olfactory signals in driving these behaviors, perhaps, indicating alternate means of integrating multi-sensory cues. *D. wrightii* produces a flower with consistent visual, morphological, and olfactory characteristics, all used as sensory cues guiding hawkmoth pollination (11, 73). In general, animal pollinated flowers tend to be conspicuous in terms of color, shape, texture, and odor gestalt (74, 75). The flower delivers a reliable multi-component signal insuring pollination success, in turn the insect relies on these signals maximizing nectar acquisition supporting metabolically costly searching behaviors (76–79). Our results indicate that innate behaviors involved in foraging – pollination based mutualistic interactions are likely dependent on a stability between the visual stimulus and the appropriate floral volatile blend. Therefore, the coordination of different senses would have a direct advantage in generating an efficient behavioral output (30, 80). In contrast to the floral stimulus multiple genetic and environmental factors qualitatively and quantitatively influence the *D. wrightii* constitutive plant foliage volatile blend, (81–83), potentially delivering variable olfactory signals. In general, leaf volatile profiles change due to the time of day (84), plant genotype (85), inter-plant communications (86), and other biotic and abiotic stress induced conditions (87–89). Insects and plants have opposing interest regarding oviposition: the insect attempts to efficiently determine the most suitable host for its offspring, while the plant aims to prevent this approach. We demonstrate that oviposition host-seeking behaviors are still functional without perceiving OR-related odorant signals from the plant. We may interpret this as an alternate behavioral approach towards integrating multi-sensory information from a host, possibly permitting greater flexibility to highly uncertain volatile blends.

In flight, odorant plumes are presented to the insect as intermittent packets of odor blends in a turbulent air stream (90) and OR-mediated olfaction is involved in directing plume tracking behaviors (91). Moth odor-guided flight behaviors are facilitated through the intermittency of odor packets in a visually aided up-wind flight path (90, 92). When analyzing the flight trajectory of *orco* KO male hawkmoths we observed a prolonged searching behavior both in time and space (Fig 4B, C’), which was followed by an abrupt alighting of the hawkmoth on the stimulus source upwind. The natural flower stimulus did not seem to be a significant cue in directing *M. sexta* flight behavior, hinting, at an innate role for combined stimulus in directing flight behaviors, as previously described in the wild hawkmoths (7).

*M. sexta* foraging is an example of a behavior that is mediated by the integration of multiple sensory inputs, combining vision and olfaction to effectively determine the salient stimulus (93, 94). Observations in wind tunnels (with 2-3m distance to the stimulus on release) or free-air fields (10-24m) with natural or semi-natural stimuli, found that presenting the visual stimulus alone would result in *M. sexta* approaching the source, but did not result in feeding (9, 95). In certain cases, only when an odor accompanied the visual stimulus an approach was followed by foraging attempts (96). Concluding that both cues are necessary for a hawkmoth to initiate and sustain foraging behavior (9, 94, 96). However, other studies using only an artificial visual stimulus, in a wind tunnel and a 1.5 m diameter circular mesh arena showed that the hawkmoth was able to extend its proboscis without a pronounced olfactory stimulus (8, 27, 97). In our study, we observe that disrupting the olfactory response towards the flower significantly ablated proboscis extension and further foraging behaviors. The *orco* KO hawkmoth exhibited a novel behavior in the presence of a natural innate stimulus: when approaching the flowering plant, they did not hover but instead landed abruptly. This behavior has not been observed in previous published reports, and we interpret this behavior to be an effect of confronting a natural stimulus with a disrupted olfactory identity. This also suggests that other stimuli, including detection of water vapor (98) and CO_2_ (71) from the flower, were not sufficient to elicit feeding. Within our behavioral paradigm we observed the interaction with the flowering plant within a small segment of time and space. It is likely that the reluctance to forage at that moment is related to the sensory and neurobiological decoupling of the olfactory system from the visual stimulus. Alternatively, while the flowering plant headspace did not elicit any responses from the antennae or AL, single odorants representative of the floral blend elicited minor responses from the ORCO KO antennae at high concentrations. In the *M. sexta* AL, distinct regions remain activated in the *orco* KO when certain odorants were used as stimiuli (Fig. 3C), indicating putative glomeruli involved in IR-mediated responses. Therefore, the hawkmoth might have a residual detection of the flower likely determined through remaining IR expressing neurons. However, this spatial pattern of activation is not representative of those eliciting proboscis extension behaviors (27). Additionally, odorants eliciting IR responses have been demonstrated to alter the valence property of that odorant in an *orco* KO background (68).

Herbivore induced plant volatiles manipulate the plant biotic environment resulting in reduced herbivory (18–20). Female hawkmoths are adapted to process olfactory information from plant odor plumes as a guide towards specific plant host. Sexually dimorphic small trichoid sensilla found on the female antennae house OSNs broadly tuned to subset of plant volatiles: terpenoids and aromatics (99), and behaviors towards relevant plant volatiles would theoretically parallel mate or flower plume seeking behaviors (100). In our experiment, gravid females that flew were able to quickly locate a *D. wrightii* non-flowering plant independent of *orco* based olfaction (Fig 5B). This observation indicates that OR-dependent olfaction might be additive to, but not crucial for this behavior, directing our attention to other chemosensory or visual cues playing a role. Our results indicate significant antennal responses from *orco* KO hawkmoths to certain odorants (Fig. 3B, Table S2). A selection of these are detected by IRs in *D. melanogaster* as well (68). IRs are an ancient chemosensory receptor gene family in arthropods (101) and IR-mediated olfaction is independent of *orco*. Among the *orco*-independent physiologically active compounds, some are released by *M. sexta* host plants, such as hexanoic acid, benzaldehyde, and farnesene (12, 102). It is likely that the M. sexta IR gene receptor family is involved in the detection of these volatiles; however, antennae and AL responses are reduced, indicating orco dependent signals contribute an additive factor to the full physiological response. It is feasible that an OR-independent olfactory input might be sufficient in guiding the female to a potential host plant. However, it should be further explored if these putative IR inputs determine the same innate behavioral valence as the full IR and OR-dependent input.

Visual perception has been attributed to contribute a large role in butterfly oviposition (103, 104). And although butterflies have a more extensive repertoire of color receptors, crepuscular or nocturnal hawkmoths are still able to employ color reception in directing their behaviors (104–106). The contrast image provided by the plant growth pattern or leaf shape may also support host searching behaviors (107). Additionally, previous studies have observed how ambient light related to the moon phases might affect *M. sexta* host localization (108). Our experiments likely indicate that a behaviorally independent visual input may play a role in host plant localization in the absence of relevant odor stimuli. Further analysis providing visual choices within the natural plant stimulus might further elucidate the extent of visual perception independent of OR-mediated signals. The gustatory sense mediates many close range Lepidoptera behaviors (109–112). A female hawkmoth tarsal contact with the plant is sufficient for oviposition, mediated through a specific pattern of tastants (110, 113, 114). Without *orco* the oviposition behavior of a gravid female was not significantly reduced, indicating that OR-dependent olfaction is non-essential under this context. In other words, while olfactory information may lead to more directed oviposition behaviors, other sensory cues might also be necessary or sufficient.

The relationship between *M. sexta* and *D. wrightii* is one of co-evolved plant chemical signals and insect chemosensory detection. Analyzing the precise contribution of each sense is informative on how insect and plants use and manipulate signals in order to enhance their overall fitness. With the *M. sexta orco* KO we observed a contextual dependency on OR-based olfaction in this relationship. In foraging, the critical nature of olfaction is likely goverened through the integration of visual and olfactory stimuli in order to elicit the complete range of foraging behavior. The coupling between vision and olfaction in this behavior highlights the evolutionary link between floral shape and color with scent for *M. sexta* pollinated flowers. On the other hand, the decoupling of OR-mediated olfaction from oviposition behaviors might favor innate adaptability towards other potential signals. In the wild this would likely provide a mechanism for female oviposition no-choice behaviors in the face of potentially unreliable environmental information. Our study presents another step in understanding the link between neurobiology and ecology through the application of CRISPR – Cas9 to classic ecological models. It allows us to speculate on a role for olfaction in the life history of a hawkmoth and how it might have adapted through the course of co-evolution. The development of feasible germ-line cell transformation will permit future studies to apply transgenic tools to further explore the neurobiology of co-evolved behaviors shaping our natural heritage and agriculture.

## Material and Methods

Procedures for M. sexta CRISPR – Cas9 sgRNA design and functional verification, *M. sexta* and *D. wrightii* general colony rearing, volatile collection and analysis, AL calcium imaging odorant stimulation, image analysis, and wind-tunnel staging and 3d-tracking are described in **Supplementary Methods**.

### *M. sexta* pre-blastoderm microinjection, embryo staging, and rearing

Eggs for microinjection were collected from a breeding chamber with a potted *D. wrightii* plant placed 15 minutes before the simulated dusk conditions. Eggs were processed for microinjection 1 – 2 hours post-oviposition. Freshly laid eggs were collected after 15 minute bouts of oviposition and stored at 5 °C until sufficient eggs were accumulated. Eggs were washed thoroughly with distilled water in a *Drosophila* egg lay cup with a mesh screen (Genesee Scientific; Flystuff.com; United States), followed by 5 minute incubation in 2 % Commassie brilliant blue stain (50 % methanol v/v and 10 % glacial acetic acid v/v). The Commassie blue treatment facilitates evaluation of *M. sexta* egg polarity based on the staining and visualization of the micropyle under a stereomicroscope (Leica S8APO; Leica; Germany), in addition to softening the egg shell for injection (115). The eggs were washed thoroughly in distilled water and dried on tissue paper before staging. Eggs were aligned on a microscope slide dressed with double-sided tape and lined up against a longwise strip of ERKOGUM or clay. Eggs were placed with posterior side relative the micropyle (anterior) facing out based on the probability of this region developing the germ plasm determinants (116). While some Lepidoptera species like *Bombyx mori*, have been shown to have an alternate method of embryonic pre-patterning, *M.sexta* has been demonstrated to follow a similar pathway to that of *Drosophila* (117, 118).

### Embryo microinjection

Eggs were injected with a short taper quartz glass capillary with filament (Sutter Instrument: ID: 1.0 mm; OD: 1.0 mm) pulled with a Sutter P-2000 laser puller (1 Line: Heat = 700; Filament = 2; Velocity = 30; Delay = 130; Pull = 75) (Sutter; United States). The underside of the glass slide with eggs was adhered with water to the glass stage plate on a Zeiss AxioZoom V16 stereomicroscope fitted with an aureka^®^ digital micromanipulator (Aura Optik; Germany) set at a 45 ° angle. A Narishige IM-300 microinjector (Narishige; Japan) with nitrogen sourced pressure standing at 62.0 psi and an initial pressure of 20.0 psi was used during initial injection with further adjustments, as needed, down to 2.0 psi. The pressure is lowered to adjust to the fine capillary tip breaking off from repeated injection, but maintains a thin enough taper to keep injecting with high survivability. After injection, wounds were sealed with cyanoacrylate adhesive (Loctite 401; Henkel; Germany) and the slide is placed in a fly vile with a wet cotton bottom and a sponge stopper, eggs were then placed in a climate chamber.

### Rearing of injected colony

On day 3 post-injection eggs were gently removed from the slide and placed on a petri dish in proximity to a small amount of larval artificial diet. Freshly hatched larvae were separated into small cups (Solo 1.25 oz Soufflé cups) with diet and sealed with custom cut cardboard lids. Larvae were transferred to a small 250 ml plastic salad boxes after reaching third instar and fed ad libitum, refreshing food when needed. Wandering stage larvae were separated into a new plastic tub with tissue paper and transferred to brown paper bags two weeks later.

### Genetic crosses

One male and one female hawkmoth were placed in a mesh screen cage (30 x 30 x 30 cm) to mate. A *D. wrightii* leaf or paper towel was placed 1-2 days later to collect eggs. Eggs were placed in a plastic cup with label of parental generation and the current progeny colony name. Once larvae hatched from the eggs, individuals were separated in to small plastic cups for genotyping.

### *M.sexta* Genotyping and Phenotyping

A 1.0 mm section of caterpillar horn from second-third instar larvae was harvested and placed in a 96-well plate with 10.0 µl of MyTaq™ Extract-PCR Kit Lysis buffer (2 µl Buffer A; 1 µl Buffer B; 7 µl ddH_2_O, Modified protocol from Bioline; Meridian Life Sciences; United States). Tissue was homogenized using a wet toothpick to quickly press the horn tissue to the side of the well or by cutting the horn multiple times in the lysis buffer. A nested PCR (Table S2) using MyTaq™ HS Red Mix (Bioline; United States) was necessary for a sufficient amplification of a 920 bp region flanking the gRNA1 target site. The nested PCR provided an approximate equivalent amplification needed for T7 Endonuclease I (T7E1, NEB; United States). The resulting digest was visualized on a 1.5 % agarose gel. PCR amplified DNA from T7E1 digest positive individuals of the G1 generation was cloned into Dual Promoter TA Cloning^®^ Kit pCR^®^II Vector (Invitrogen; ThermoFisher Scientific; United States). Multiple colonies were picked for each cloning reaction, amplified and purified in a 96 well plate and sequenced at the Max Planck Institute for Chemical Ecology. Sequenced data was analyzed using Geneious version 8 (Biomatters; United States). Three sequences hence 3 colonies with positive indel mutations were confirmed by sequencing (Fig. S1), once these individuals were backcrossed to WT male *M. sexta* the rest were disposed of by freezing. Minimal colonies were maintained because of space limitations in the climate chamber. Position of this frameshift mutation was identified using TOPCONS2 to predict the *M.sexta* Orco topology and illustrated using Protter (*http://wlab.ethz.ch/protter*) (Figure 1E). Once the G2 generation of colony 1 was confirmed by T7E1 the other colonies were discarded.

The stochastic insertion of a BclI site in colony 1 facilitates the identification of heterozygous (HET) and knock-out (KO) individuals from the WT individuals. To maintain the following generations for experimental purposes primers were designed to amplify 201 bp (Table S3) with the target site or mutation located 50 bp downstream from the 5’ of the PCR amplicon. A 10 µl BclI digest (5 µl PCR template, 1 µl NEB3.1 Buffer, 0.25 BclI, 3.75 ddH_2_O) reaction was set up from the PCR reaction and loaded on a 2 % agarose gel. Every subsequent generation requires identification of HET and KO individuals using BclI digest. Accuracy of genotyping method was examined by electroantennogram (EAG) characterization of individual hawkmoths after behavioral experimentation. Either the full panel of odorants (Fig. 2B) or a selection of five diagnostic odors (Bombykal, Hexanoic Acid, Linalool, Pyrrolidine, Benzyl Alcohol) was used to match antennal responses to genotyped individuals.

## Supporting information

Suporting Information (SI)

