## Supplementary material for "Mutagenesis of *orco* impairs foraging but not oviposition in the hawkmoth *Manduca sexta*": Suporting Information (SI)

**Figure S1**

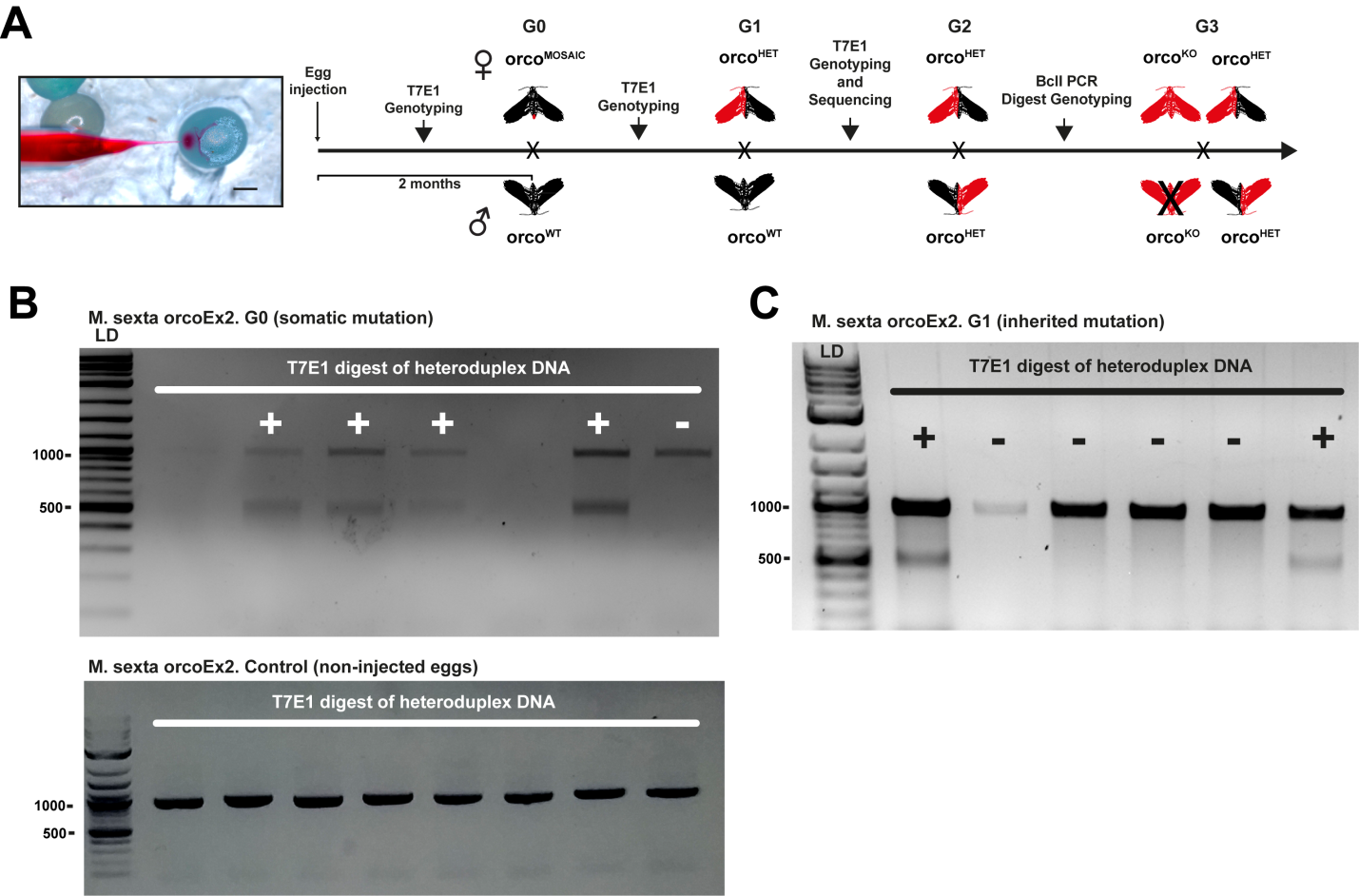

**Figure S1*. M. sexta* injections and initial mutation detection.** (A) With a specially designed quartz glass microcapillary CRISPR-Cas9 RNP mixture diluted in Acid Red 27 was injected into a *M. sexta* egg (Scale bar: 0.5 mm). *M. sexta* has a generation time of 2 months. Sequencing and T7 Endonuclease I (T7E1) genotyping was used to identify CRISPR-Cas9 efficiency in the G0 and inherited mutations in the G1. Hawkmoths were crossed back with WT in the G1 to stabilize the mutation 1 with BclI restriction enzyme site in the KO allele. Subsequent genotyping was performed for G2 and onwards for every experimental animal, since *orco* KO *M. sexta* males do not mate (indicated by an X). Thus, a stable line could not be established. KO = homozygous mutants with null mutation of orco, HET = heterozygous hawkmoths. (B, top) T7E1 digest of PCR product from injected individuals (G0). The digest is indicative of a mutated allele (+) when forming the heteroduplex of the PCR product at ~ 500 bp. (B, bottom) Control experiment of non-injected individuals. The T7E1 digest of heteroduplex DNA in the mutation detection assay shows no evidence of cuts. (C) T7E1 digest of PCR product from G1 individuals. Positive sign indicates germ-line cell inherited mutations found in the G1 generation.

**Figure S2**

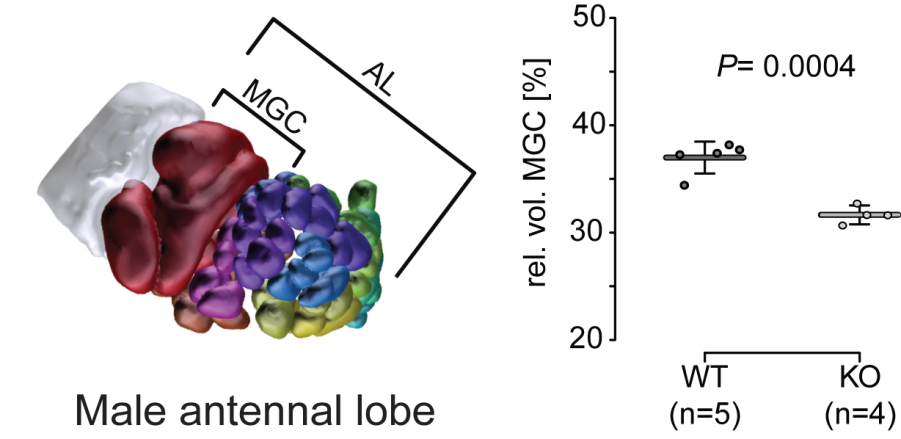

**Figure S2. Relative volume of the male macro-glomerular complex (MGC) is reduced in the *orco* KO.** Left side shows reconstruction of the male antennal lobe (AL) with the MGC highlighted in dark red. Right side shows mean relative volumes (horizontal bar) and standard deviation (error bars). Filled circles represent individual measurements. Welch corrected t-test was used to compare treatments.

**Figure S3**

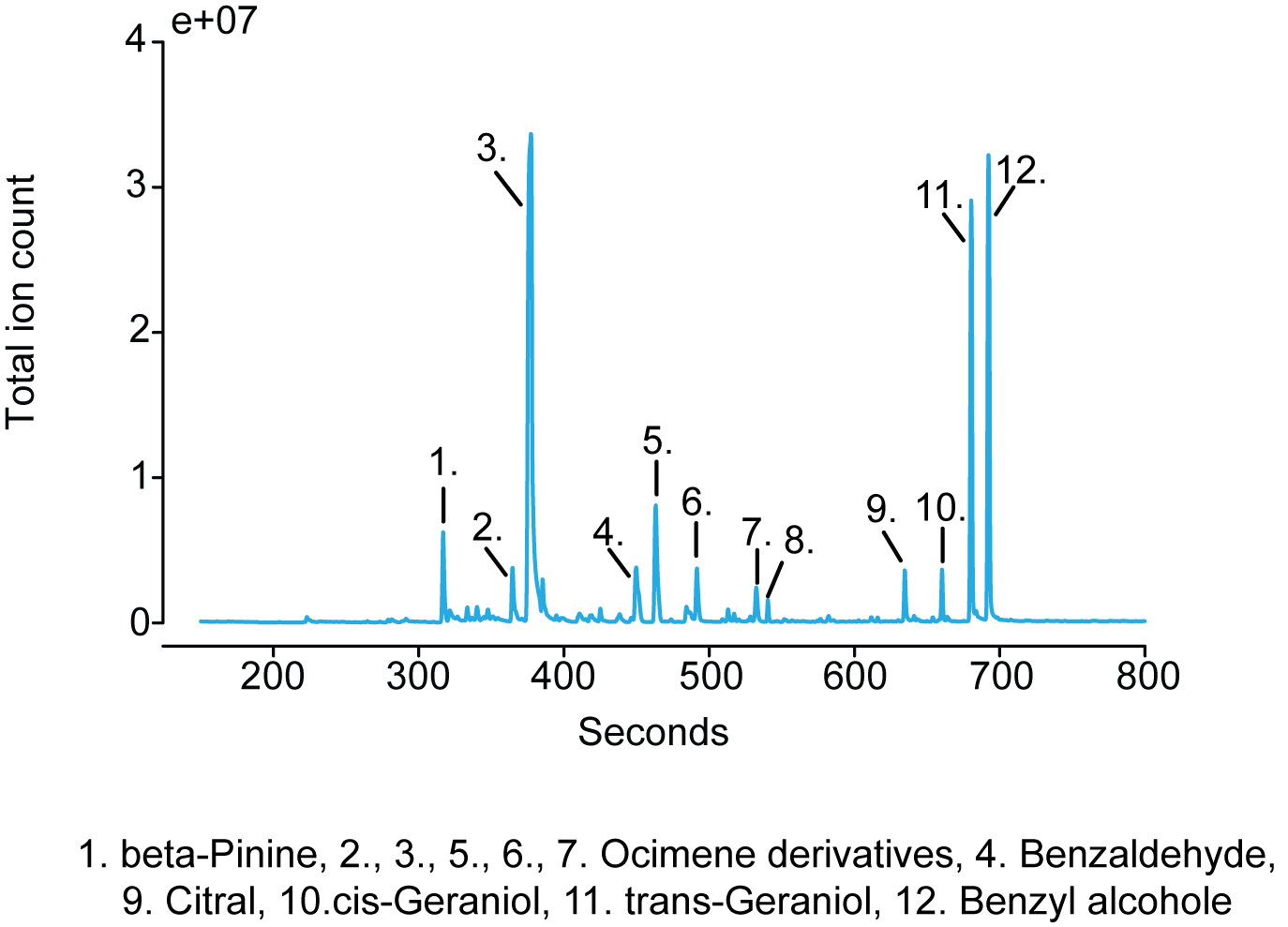

**Figure S3. Chromatogram of the headspace of a flowering *D. wrightii* plant.** Odorants denoted by the numbers were identified by comparison with the NIST 2.0 library.

**Figure S4**

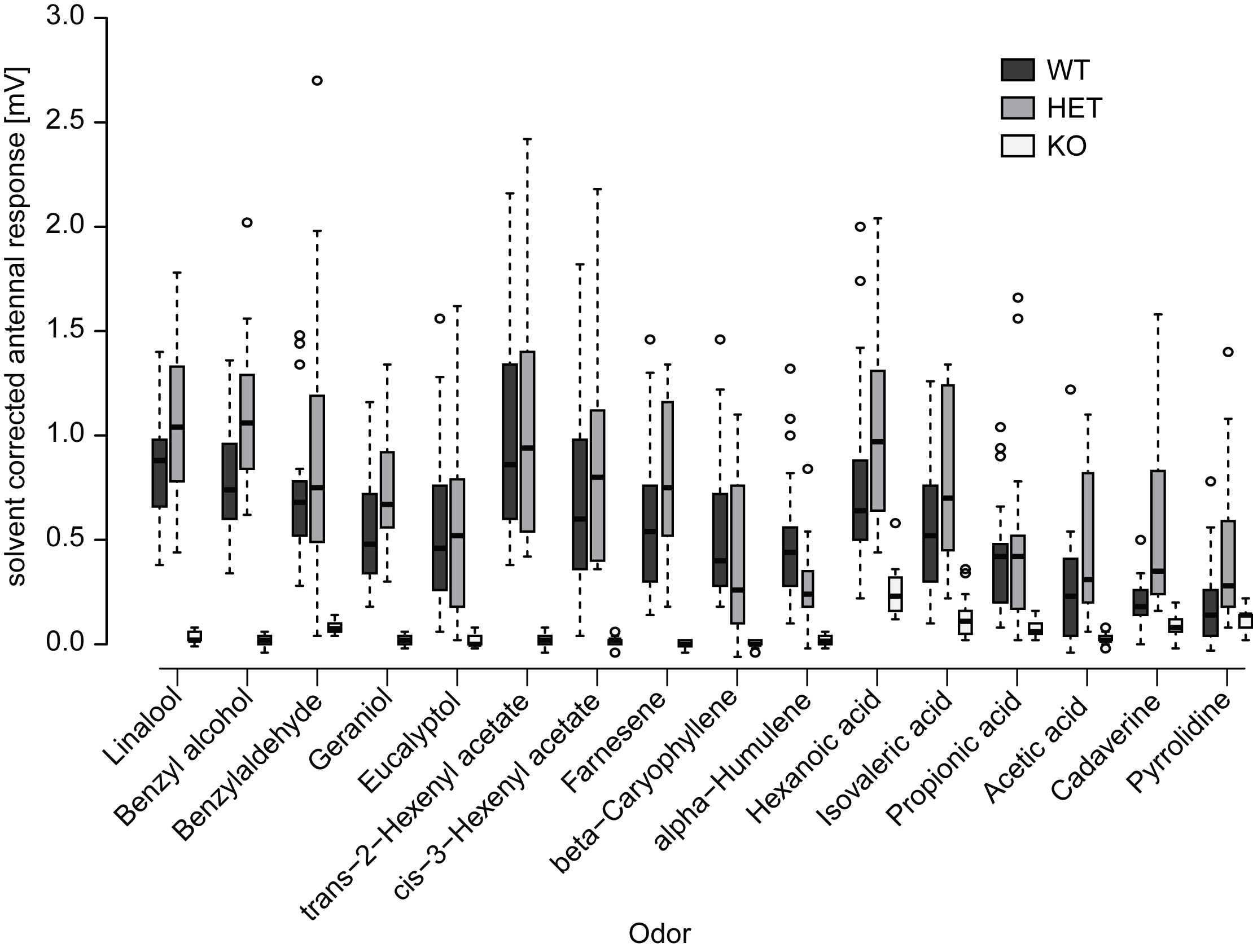

**Figure S4. Male EAG responses to ecologically relevant volatiles.** For each genotype, 18 (KO) or 21 (WT and HET) animals were tested. Significance levels are shown in Table S1.. Boxplots show the median, solvent corrected EAG amplitude (horizontal line in the box), the 25^th^ and 75^th^ percentiles (lower and upper margins of the box) together with the 1.5 x interquartile range (whiskers), and points laying outside the 1.5 x interquartile range (circles).

**Table S1.** Median values, interquartile ranges and significance levels of male antennal responses for WT (n=21), HET (n=21), and KO (n=18) moths to different odorants as well as significance levels for differences between the three genotypes in response to the individual odorants. Significances of the responses of each genotype to the individual odorants were tested by comparing the corrected response values against zero using a Wilcoxon rank-sum test. Significance levels between the different genotypes were determined by comparing the corrected response values towards the different odorants using a Kruskal-Wallis test, followed by a Holm-corrected Wilcoxon rank-sum test.

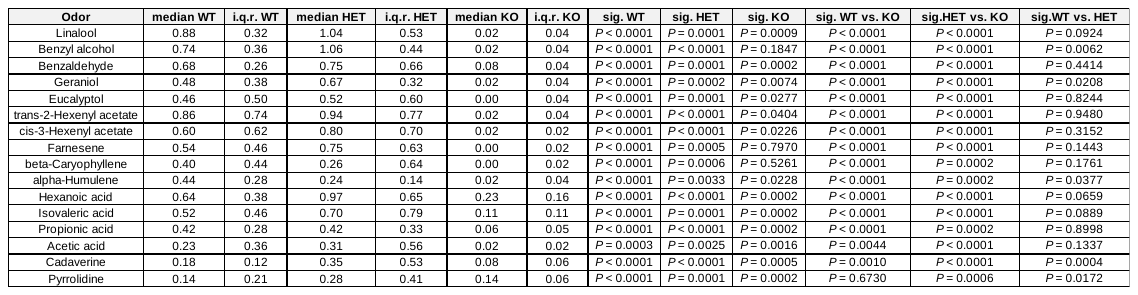

**Figure S5**

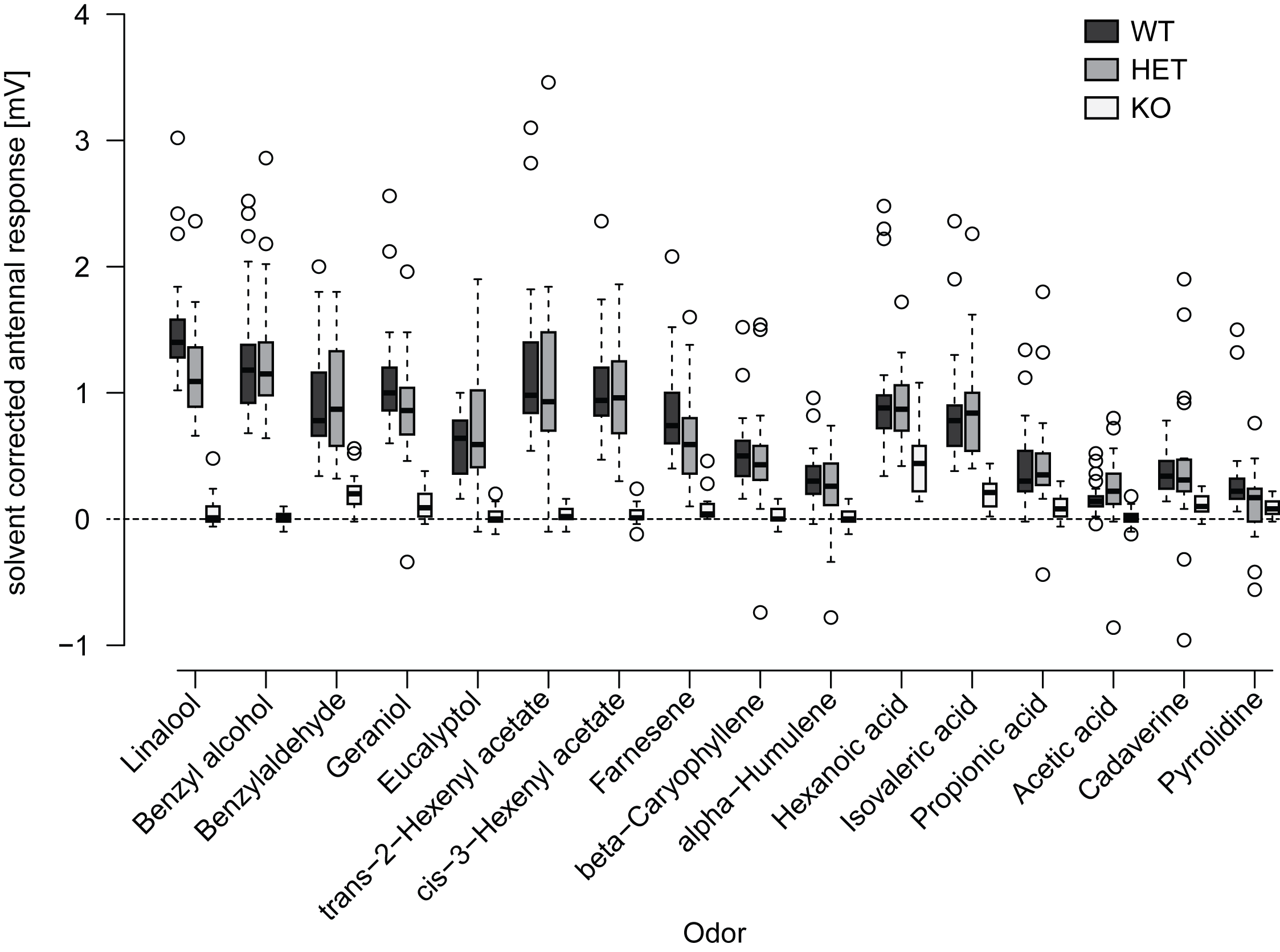

**Figure S5. Female EAG responses to different ecologically relevant volatiles.** For each genotype, 18 (KO), 20 (HET) or 21 (WT) moths were tested. Significance levels are shown in table S2. Boxplots show the median, solvent corrected EAG amplitude (horizontal line in the box), the 25^th^ and 75^th^ percentiles (lower and upper margins of the box) together with the 1.5 x interquartile range (whiskers), and points laying outside the 1.5 x interquartile range (circles).

**Table S2.** Median values, interquartile ranges and significance levels of female antennal responses for WT (n=21), HET (n=20), and KO (n=18) moths to different odorants as well as significance levels for differences between the three genotypes in response to the individual odorants. Significances of the responses of each genotype to the individual odorants were tested by comparing the solvent corrected response values against zero using a Wilcoxon rank-sum test. Significance levels between the different genotypes were determined by comparing the corrected response values towards the different odorants using a Kruskal-Wallis test, followed by a Holm-corrected Wilcoxon rank-sum test.

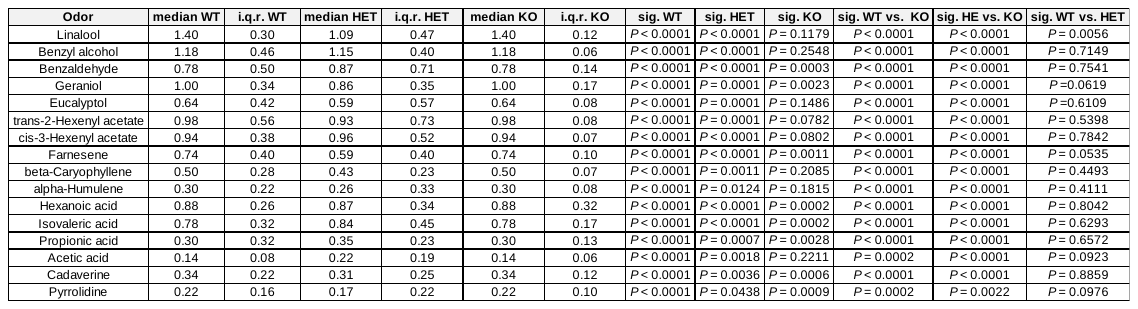

**Figure S6**

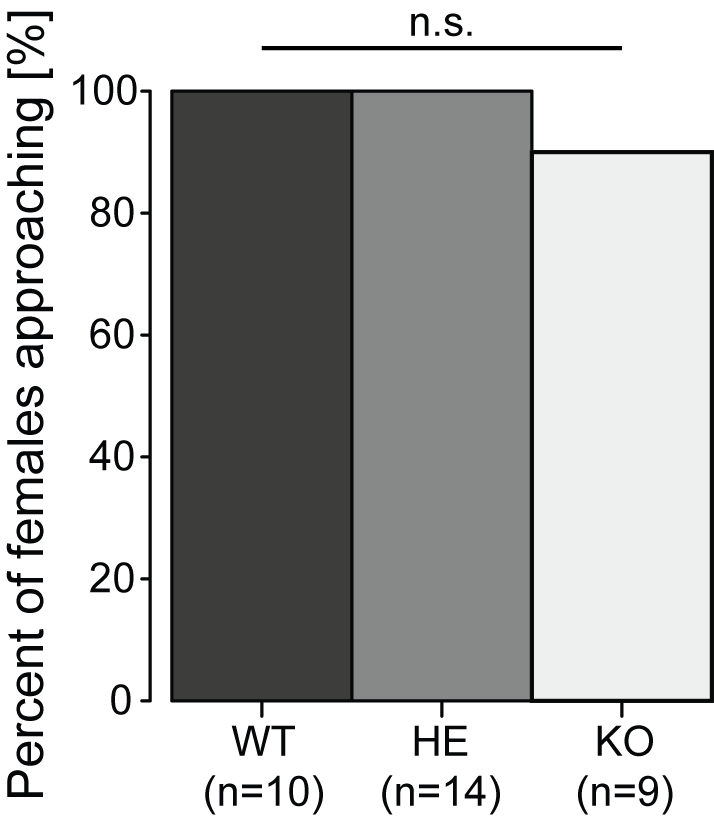

**Figure S6. Percentage of virgin female moths approaching flowers after taking flight.** Significance levels between WT, HET and KO were tested using Holm-corrected Fisher’s Exact tests**.**

**Figure S7**

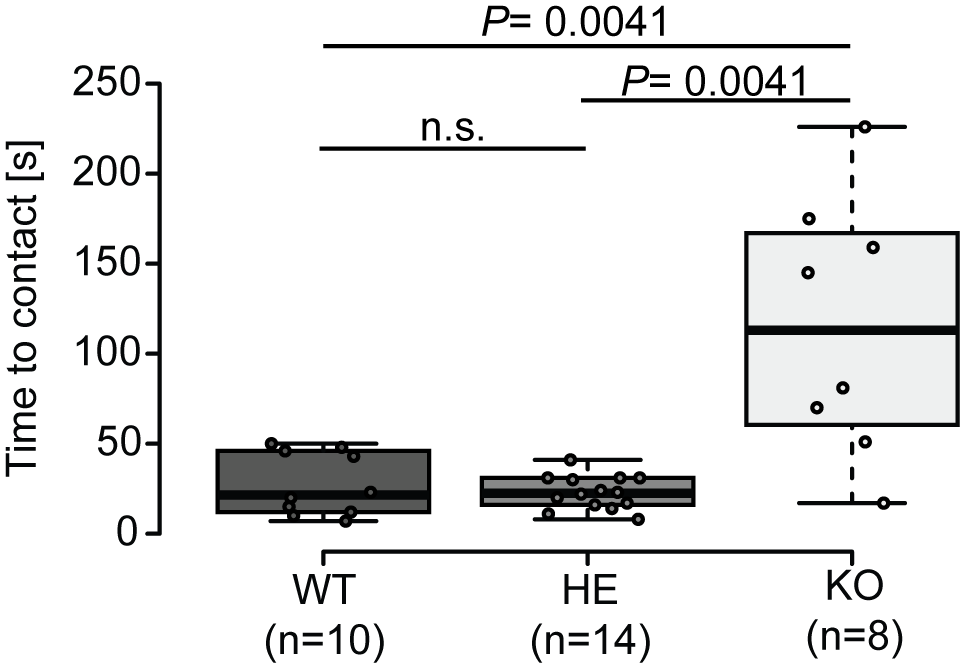

**Figure S7. Time from take-off to first flower contact for virgin female moths.** Differences between genotypes were tested using a Kruskal-Wallis tested followed by a holm-corrected Wilcoxon rank-sum test. Boxplots show the median time to contact (horizontal line in the box), the 25^th^ and 75^th^ percentiles (lower and upper margins of the box) together with the 1.5 x interquartile range (whiskers). Points indicate individual measurements.

**Figure S8**

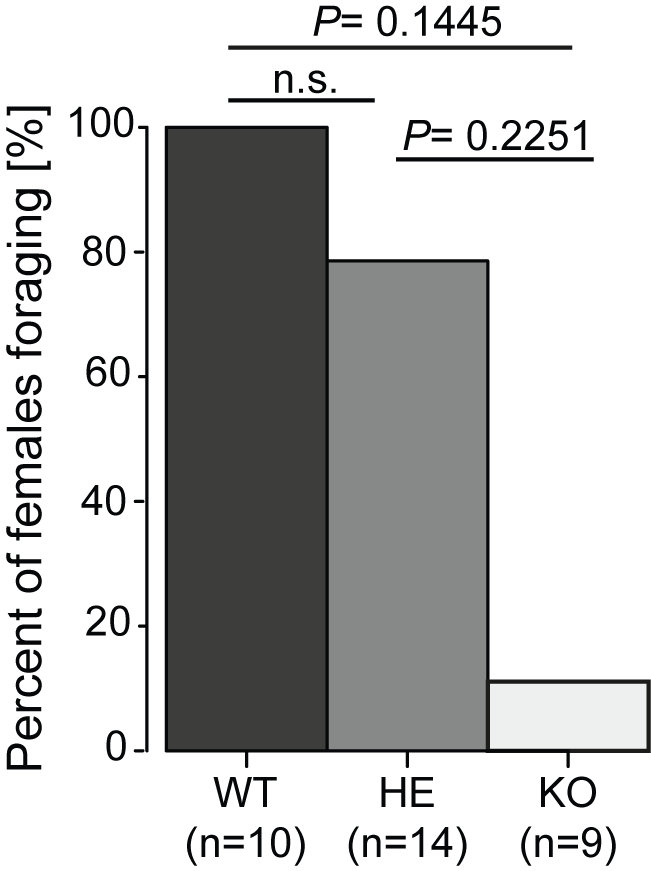

**Figure S8. Percentage of virgin female moths foraging after approaching the *D. wrightii* plant.** Significance levels between WT, HET and KO were tested using holm-corrected Fisher’s Exact tests**.**

**Supplementary Methods.**

**Wild-type *M.sexta* rearing**: **General rearing practice.** Wild-type hawkmoths were placed in a breeding chamber consisting of a large screen cage 76 x 42 x 122 cm (Exoterra, Hagen, Canada) with a potted *D. wrightii* plant and allowed to freely mate and oviposit. The breeding chamber was placed within a climate chamber under long day light cycle 16 h: 8 h light: dark, 27: 25°C cycle at 30 % humidity. Eggs were hatched in close proximity to artificial diet in 250 ml plastic salad boxes. Hatched larvae were housed in food storage plastic boxes and fed ad libitum, avoiding humidity, until wandering stage where they were placed in wooden blocks to pupate. After 2 weeks pupae were taken out of the wooden blocks, sexed, and placed in male or female only climate chambers 16 h: 8 h light: dark, 27: 25°C cycle at 70 % humidity.

**Plant material.** *Datura wrightii* plants were grown at the Max Planck Institute for Chemical Ecology, Jena, Germany, from seeds originally collected at D.I. Ranch, Santa Clara, Utah, USA. Directly after potting, young plants were transferred to a York Chamber (Johnston Controls, USA) with the same light regime and climatic conditions as the hawkmoth flight cages. Plants were watered daily with 100 ml tap water supplemented with 0.12 g / L fertilizer (Peters Professional Allrounder, Planta Duengemittel, Germany, nutrient composition: 20% N, 20% P2O2, 20% K2O, 0.015% Cu, 0.12% Fe, 0.06% Mn, 0.01% Mo and 0.015% Zn). For oviposition experiments, plants were used at a stage in which the first flower buds had just emerged, with each plant being only used once. For pollination experiments plants were used as soon as the first flower opened. In those cases were several flowers were present at the same stage, all but one flower was removed on the day before the experiment.

**Volatile collection and analyses.** To collect the headspace of flowering *D. wrightii*, plants were enclosed in a glass cylinder (49 × Ø34 cm). The cylinder was closed-off just above the pot using a movable plastic (polyoxymethylene) door with an opening tightly fitting the plant stem. Charcoal filtered air was pushed into the cylinder at a rate of 0.5 L/min, while air was simultaneously pulled from the cylinder at 0.4 L/min through a glass tube (ARS, USA); packed with glass wool and 20 mg of Super Q (Alltech, Germany). All odor collections were performed for 8 h from the beginning until the end of the scotophase in a York chamber at 25º C and 70% r. H. Thereafter the SuperQ filters were eluted with 400 μl dichlormethane and stored at -20º C until further use. ***Chemical analysis***. 1 μl of the sample was injected into a gas chromatograph-coupled mass spectrometer (GC-MS) (Agilent 6890 GC & 5975C MS, Agilent, USA). The gas chromatograph was equipped with a polar Innowax column (30 m, 0.25-mm ID, 0.25-µm film thickness; J&W Scientific). The temperature of the inlet port was maintained 240 °C and GC oven was set to an initial temperature of 50 °C, which was held for 2 min. Thereafter, the temperature was increased at a rate of 13 °C/min to 250 °C, which was again held for 5min. The MS transfer line was held at 280 °C and the MS operated in electron impact mode (70 eV, ion source: 230 °C, quadrupole: 150 °C, mass scan range: 33–350 m/z, scanning rate 4.42 scan / s). Compounds were identified by comparing mass spectra against the NIST 2.0 library.

**Immunohistochemistry and antennal lobe measurements.** An Orco KO phenotype has previously been shown in some insect species to cause defects in AL morphology (1, 2). We investigated if any such effect is observed in the *M. sexta* brain using immunohistochemistry and virtual brain reconstructions, similar to what has been previously described (3, 4). Brains of 3 day old male hawkmoths were dissected in phosphate-buffered saline (PBS) after the hawkmoth was tested by EAG. The tissue was then fixed overnight at 4º C in a solution of one part formaldehyde (37%, Roth, Germany), one part methanol, and eight parts PBS. Thereafter brains were rinsed in PBS for 1h and pre-incubated in 5% normal goat serum (NGS) (Jackson ImmunoResearch, United States) in PBS containing 0.3% Triton X-100 (PBS-T) (Sigma-Aldrich, United States). Next a primary mouse anti- SYNORF-1 antibody (DSHB) against a fusion protein consisting of a glutathione-S-transferase and the first amino acids encoded by 5’- end of the *Syn* 1 gene (5), which previously been shown to reliably stain neuropils in *M. sexta* (3), was applied at a ratio of 1:100 in 1% NGS diluted in PBS-T. . Brains were incubated in this solution for 6 days at 4º C. Afterwards the samples were rinsed five times for 20 min in PBS-T before the secondary goat anti-mouse antibody linked to Alexa Flour 488 (Life technologies, United States) was applied at a ratio of 1:300 in 1% NGS-PBS-T. Brains remained in this solution for 4 days at 4º C. The secondary antibody was then washed of by rinsing the samples six times for 20 min in PBS-T. Finally, brains were dehydrated in an increasing ethanol series (50%, 70%, 80%, 90% and two times 100%, with 15 min each) and cleared in methyl salicylate overnight. The samples were scanned using a multiple-photon confocal laser scanning microscopy (MPCLSM) (Zeiss laser scanning microscopy [LSM] 710 NLO confocal microscope; Carl Zeiss, Germany). All preparations were viewed under a 20x air objective (Plan-Apochromat 20x/0.8 M27; Carl Zeiss, Germany) in combination with the internal Argon 488 (LASOS) laser. Antennal lobes were reconstructed manually using the software AMIRA version 5.5.0 (FEI Visualization Sciences Group). The antennal lobe of male hawkmoths is characterized by several enlarged glomeruli called, macro-glomerular complex (MGC), which receives its input from pheromone sensitive neurons on the antenna (6).

***M.sexta* *orco* KO Functional Characterization. *Electro-antennograms (EAGs).*** Electro-antennograms were used to determine the stimulus-dependent potential changes summed over the whole hawkmoth antenna. Different ecologically relevant odorants were tested and EAG responses of the different genotypes the *orco* KO, *orco* HET, and WT hawkmoths were compared. For this we removed one antenna of the hawkmoths just above the scapulum and immediately placed the cut end into a glass capillary containing *M. sexta* ringer solution (150 mM NaCl, 3 mM CaCl_2_, 3 mM KCl, 10 mM TES buffer, 25 mM Sucrose). In order to increase conductivity, the tip of the antenna was cut after the third flagellum and placed into a capillary containing the same saline solution. Both capillaries were then placed on Ag-AgCl wires, of which one was connected to a grounding electrode, while the other was connected to a 10x a high-impedance d.c. amplifier (Syntech; The Netherlands). The signal was then sent to an analog/digital converted (IDAC-4, USB, Syntech, The Netherlands) and transferred to a PC. Finally, the data were analyzed and saved using the software package Autospike v3.2 (Syntech; The Netherlands). All hawkmoths tested in the EAGs, were genotyped before as larvae, had also been used in the wind tunnel tests, and were 3-4 days old. Odorants for EAG analyses were selected based on a literature review as well as on compounds identified in the headspace of flowering *D. wrightii* plants. The main pheromone compound of *M. sexta*, bombykal, was kindly provided by Prof. Dr. Aleš Svatoš and Dr. Jerrit Weißflog from the Mass Spectrometry Group at the Max Planck Institute for Chemical Ecology, Jena, Germany. In order to stimulate the antenna with a certain odorant 10 μl of the odor at a dilution of 10^-2^ was pipetted onto a filter paper (Ø 12 mm, Whatman, Sigma-Aldrich; United States) and placed into a glass pipette. The pipette was then connected to a stimulus controller (Syntech Stimulus Controller CS-55), which if triggered, alternated the air flow of 0.4 l/min with a parallel clean air stream, delivering an odor pulse of 0.5 s. This pulse pushed the odorant from the pipette into a humidified air stream running at a constant speed of 0.8 l/min. This air stream finally delivered the odorant to the hawkmoth antenna.

***Single sensillum recordings (SSR).*** The response of the hawkmoth to bombykal was investigated by recording the physiological response of individual pheromone-sensitive trichoid sensilla. For this, three day old naïve male hawkmoths were immobilized in pipette tips with the antenna fixed on a microscope slide using dental wax (Boxing wax, Sybron/Kerr; United States). The apical segments of the antennae were removed with micro-scissors. A glass electrode filled with hemolymph Ringer was used as indifferent electrode and was inserted in the truncated end of the male’s antenna (7). The recording electrode filled with sensillum lymph Ringer was slipped over the truncated sensillum. Signals were amplified (10x a high-impedance d.c. amplifier, Syntech, The Netherlands), sampled (10,667 samples/s), and filtered (100–3000 Hz with 50/60 Hz suppression) via an analog/digital converter (IDAC4, Syntech, The Netherlands) connected to a computer. Action potentials were identified using Syntech Auto Spike 32 software. For odor delivery a system similar to the EAG set-up was used.

**Antennal lobe functional imaging analysis*.*** Hawkmoths were pushed in a 15 ml plastic tube with the tip cut open. The head was protruding at the narrow end and was fixed in this position with dental wax. Labial palps and proboscis were also fixed with wax to reduce movement artefacts during the experiments. A window was cut in the head capsule between the compound eyes and tissue covering the brain was removed until the antennal lobes were visible. The membrane-permeable form of a fluorescent calcium indicator (Calcium Green-1 AM, Invitrogen) was dissolved in physiological saline solution physiological saline solution to remove excessive dye, and hawkmoths were stored at 4°C over night. Imaging experiments were performed the following day. The imaging set-up consisted of a CCD camera (Olympus U-CMAD3) mounted to an upright microscope (Olympus BX51WI) equipped with a water immersion objective (Olympus, 10x/0.30). Calcium green-1 AM was excited at 475 nm (500 nm SP; xenon arc lamp, Polychrome V, Till Photonics) and fluorescence was detected at 490/515 nm (DCLP/LP). The set-up was controlled by the software Tillvision 4.6 (Till Photonics). Four-fold symmetrical binning resulted in image sizes of 344 x 260 pixels with one pixel corresponding to an area of 4 µm x 4 µm.

***Odorant stimulation*.** We tested 3 monomolecular odorants (bombykal, benzaldehyde, hexanoic acid) diluted in mineral oil, and the headspace of a *D. wrightii* flower dissolved in dichloromethane. Ten µl of a stimulus were applied onto a circular piece of filter paper (diameter: 12 mm, Whatman); 10 µl of solvent served as a control stimulus. Filter papers were inserted into glass pipettes and were renewed every day. The immobilized hawkmoth was placed upright under the microscope. A glass tube was directed to one antenna (diameter: 5 mm; ending 10 to 15 mm from the tip of the antenna) delivering a constant stream of clean moistened air (0.1 l/min). Two glass pipettes were inserted through small holes in the tube. One pipette (inserted 5.5 cm from end of tube) was empty and added clean air to the continuous airstream (0.5 l/min). This airstream could automatically be switched (Syntech Stimulus Controller CS-55) to the second pipette (inserted 3.5 cm from end of tube) that contained an odorant-laden filter paper. By this procedure the airstream reaching the antenna was not altered during odorant stimulation, thus reducing mechanical disturbances. One odorant stimulation experiment lasted 10 s and was recorded with a sampling rate of 4 Hz corresponding to 40 frames. The time course of an odorant stimulation experiment was as follows: 2 s clean airstream (frame 1 to 8), 2 s odorant airstream (frame 9 to 16), and 6 s clean airstream (frame 17 to 40). Odorants were presented with at least 1 min interstimulus interval to avoid adaptation. The sequence of stimulations changed from animal to animal. Stimulations with each of the two solvents were presented twice to each animal.

***Processing of calcium imaging data***. Stimulation experiments resulted in a series of 40 consecutive frames that were analysed with custom written software (IDL, ITT Visual Informations Solutions). Several processing steps were applied to enhance the signal-to-noise ratio: i) background correction: background activity was defined as the average fluorescence (F) of frames 3 to 7, i.e. before stimulus onset, and was subtracted from the fluorescence of each frame. This background-corrected value (ΔF) was divided by the background fluorescence to get the relative changes of fluorescence over background fluorescence for each frame (ΔF / F); ii) bleaching correction: the fluorescent dye is bleaching slowly during the exposure to light. Therefore we subtracted from each frame an exponential decay curve that was estimated from the bleaching course of frame 3 to 7 and frame 26 to 40, i.e. just before and after stimulus and response; iii) median filtering: a spatial median filter with a width of 7 pixels was applied to remove outliers; iv) movement correction: possible shifts of the antennal lobe from one stimulation experiment to the next one were corrected by aligning frame 20 of each experiment to frame 20 of the median experiment in a given animal. The outline of the antennal lobe and remains of tracheae served as guides for this movement correction procedure.

***M. sexta* *orco* KO *wind tunnel no-choice assay*.** In the present study the consequences of a loss of *orco* on both foraging and oviposition were tested to examine importance of this gene mediating these ecological behaviors. Pupae from genotyped individuals were sexed, placed in paper bags, and separated into male and female specific chambers approximately one week pre-eclosion. Emerged male hawkmoths were collected for behavioral experiments 3 days post-eclosion. Gravid females were collected 1 – 2 days post mating. All animals were starved from eclosion until the experiments. On each experimental day either male or female hawkmoths were tested. At first, hawkmoths were acclimatized at 25 º C and 75% relative humidity for 1 hour before the experiments in a separated pre-incubation chamber. Then, the animals were introduced individually onto a small platform at the downwind-end of the wind tunnel and gently touched to initiate wing fanning. Based on previous studies, the conditions in the wind tunnel were set to 25 ºC and 75% relative humidity, a wind speed of 0.4 m/s and a light level of 0.5 lux (8). Hawkmoths were then allowed to fly freely in the wind tunnel for 5 minutes. During this time the behavior of the animals was observed using a custom build 3d tracking system (9). For this the animal was recorded by 4 USB cameras (Logitech C615; United States, infrared filter) at 30 Hz and a resolution of 800 × 600 pixels (each pixel corresponding to 0.3 cm^2^). The 3d position of the hawkmoth was then calculated at a rate of 10 Hz based on a background-subtraction method implanted in C., further analyses of the data was then performed in R (10). In addition to this, a fifth camera (Logitech C615; United Sates, infrared filter) had been positioned at the upwind end of the wind tunnel to closely observe the behavior of the hawkmoth on the flower or plant.

Male and virgin female hawkmoths were offered a *D. wrightii* plant with a single freshly opened flower. The number of hawkmoths foraging, defined as contacting the flower with the proboscis, as well as the time to first contact were recorded by direct observation through the close-up camera. Gravid female hawkmoths were offered a non-flowering *D. wrightii* plant, which was positioned 30 cm from the upwind end, at the center of the wind tunnel. Time to first contact and abdomen curling were noted from video recording. Eggs laid on the plant were collected, counted and stored in the climate chamber. Data from mated females was only included in the further analyses only if the mating status of the female was confirmed through egg hatching.

***M. sexta* CRISPR- Cas9 sgRNA design and functional verification*. In silico*.** The *M. sexta* genome version 1.0 was accessed through the i5k Workspace@NAL (i5k.nal.usda.gov) and imported into Geneious version 8 (11). Gene feature file (.gff) for the official gene set 2 (OGS2.0) was imported and mapped onto the genome, providing the annotated features and predicted exon - intron junctions of annotated genes (12, 13). The *M. sexta* OGS 2.0 names for olfactory receptor genes were accessed through a previous publication by Koenig et.al. 2015 and used to identify the correct genes on the genome scaffold (13). The *M. sexta* gene sequence was blasted against the genome to locate the gene sequence within a scaffold or the OGS2.0 name was used in the *M. sexta* webApollo to locate scaffold and position of the gene. The *M. sexta* genome v.1.0 (Mansexv1.0) fasta file and the GFF3 file were submitted to the CHOPCHOP ([http://chopchop.cbu.uib.no](http://chopchop.cbu.uib.no/)) database for CRISPR-Cas9 target selection sites and off-target searches (14, 15). The OGS2.0 gene names, i.e. Msex2.12779-RB, were selected to access the web tool selection of target site for *M. sexta* genes. The CHOPCHOP web tool was set to search both strands for CRISPR guide RNA (gRNA) target sites using the 5’ – N_20_NGG – 3’ motif, providing target sites for all coding regions based on GFF annotations in addition to cross-referencing the genome for potential off-target sites. The *M. sexta orco* gRNA1 and 2 were selected based on no detectable off-target sites with gRNA 1 containing only 1 mismatch, which is in the acceptable range for predicted gRNA binding efficiency (16). ***In vitro***. Recombinant *Streptococcus pyogenes* Cas9 protein with 6 – His tag and nuclear localization signal (NLS) from SV40 (PNA Bio Inc) and sgRNAs were synthesized via T7 in-vitro transcription (IVT) from fill-in PCR amplicons (17). Cutting efficiency of sgRNAs was confirmed by in-vitro digest containing PCR amplicon of target site from genomic DNA, Cas9 peptide and nuclease buffer from NewEngland Biolabs and IVT generated sgRNAs. ***In vivo***. A final concentration of 250 ng / μl sgRNA, 200 ng / μl of Cas9 peptide was added to 10 μl of dH_2_O mixed with 0.05 % of Amaranth dye - Acid Red 27 (Sigma-Aldrich) and 1.0 – 2.0 μl of solution was loaded onto quartz glass microcapillaries with a microloader pipette tip (Eppendorf). Evaluation of the injected individuals G0 mosaic and G1 generation did not reveal any large insertion or deletion events through visual inspection of PCR product on a 1.5 % agarose gel. Sequencing of individuals from G1 showed three different mutations all from the sgRNA1 sequence (Fig. S1B). The CHOPCHOP v2 sgRNA target site efficiency score ranges from 0.00 to 1.00 with 0.69 being the highest score for *M. sexta* ORCO targets, sgRNA1 and sgRNA2 are given a score of 0.47 and 0.45 respectively. While an *in vitro* digestion assay revealed that both sgRNAs were fully functional it is likely that sgRNA2 did not function *in vivo*, possibly obstructing any large indel formation.

**Supplementary References.**

1.	Yan H, Opachaloemphan C, Mancini G, Cell YH (2017) An Engineered orco Mutation Produces Aberrant Social Behavior and Defective Neural Development in Ants. *An Eng orco Mutat Prod Aberrant Soc Behav Defective Neural Dev Ants*.

2.	Trible W, Olivos-Cisneros L, McKenzie SK, Cell SJ (2017) orco Mutagenesis Causes Loss of Antennal Lobe Glomeruli and Impaired Social Behavior in Ants. *orco Mutagen Causes Loss Antennal Lobe Glomeruli Impair Soc Behav Ants*.

3.	El Jundi B, Huetteroth W, Kurylas AE, Schachtner J (2009) Anisometric brain dimorphism revisited: Implementation of a volumetric 3D standard brain in *Manduca sexta*. *J Comp Neurol* 517(2):210–25.

4.	Homberg U, Montague RA, Hildebrand JG (1988) Anatomy of antenno-cerebral pathways in the brain of the sphinx moth Manduca sexta. *Anat antenno-cerebral pathways brain sphinx moth Manduca sexta*.

10.	R Core Team (2017) R: A Language and Environment for Statistical Computing.
